# Transposon-driven pervasive genome restructuring and regulatory evolution in early-diverging Holomycota

**DOI:** 10.64898/2026.11.20.768218

**Authors:** Carlos Lax, Ghizlane Tahiri, Daniel Croll

## Abstract

Early-diverging non-Dikarya fungi showcase major evolutionary transitions in the fungal kingdom, share features with other eukaryotic clades and bacteria, and include pathogens of significant concern. These lineages contain some of the most reduced and expanded genome architectures of the fungal kingdom and have adopted unique DNA methylation systems. Yet, the evolutionary trajectory of early-diverging non-Dikarya fungi remains poorly understood. A key driver is thought to be transposable elements, which shape genomic plasticity, organization and regulation. Here, we perform a comprehensive comparative analysis across the Holomycota to chart drivers of early fungal evolution. We demonstrate that contrasting transposon dynamics directly shape genomic diversity across these lineages, fueling extensive protein domain expansions through widespread co-option. This heterogeneous landscape is regulated by a multi-layered epigenetic network impacting chromatin configuration, and we reveal a distinct transcriptional activation role for adenine methylation over specific transposons. Beyond driving functional innovation, mobile elements serve as a source of regulatory diversification, providing transcription factor binding sites at key chromosomal features. Furthermore, synteny tracking uncovers multiple whole-genome duplication events across Mucoromycotina, where DNA transposons preferentially accumulate at duplication breakpoints and undergo rapid inactivation. Strikingly, we also show that horizontal transposon transfers frequently cross deep evolutionary boundaries, including cross-kingdom exchanges with protists. Together, our findings illuminate key drivers of genome architecture in non-Dikarya fungi and reshape our understanding of genome dynamics across the fungal kingdom.

## INTRODUCTION

Fungi are ecologically highly successful and exhibit exceptional diversity and distribution across Earth’s ecosystems^1^. These ubiquitous organisms are also highly relevant due to their applications in food and antibiotic production, and their pathogenic potential on a broad range of plant and animal hosts, including amphibians and humans^2^. Our understanding of the fungal kingdom is severely limited, as most model species share either a haploid-dominant life cycle, non-motile filamentous growth with septation, mushroom production or yeast forms. Furthermore, our understanding of fungal biology is often limited to terrestrial lifestyles. This narrow scope is defined by the dominant features of the subkingdom of Dikarya^3–5^. However, from the Opisthokont ancestor shared by Metazoa and Fungi, several clades diverged earlier than the Dikarya branch. These clades are conventionally grouped under the subkingdom known as early-diverging fungi (EDF) or non-Dikarya fungi (NDF)^5^. Importantly, these clades significantly expand the diversity of traits associated with the fungal kingdom, revealing striking similarities with Metazoa, such as the presence of flagella^6^ and membrane-bound cholesterol^7^, the use of cobalamin^8^ or the retention of E2F-like proteins for cell cycle control^9^. The recognition of this shared ancestry with Metazoa had a profound impact on the reconstruction of the last universal fungal ancestor^10–13^.

The taxonomic organization and phylogenetic positioning of NDF lineages are under debate, but up to nine distinct phyla have been proposed. Together with Dikarya and its sister lineages of Cryptomycota, Aphelids, Microsporidia and Nucleariida, they constitute Holomycota^14–16^. While representatives of the Dikarya subkingdom are dominant in most ecosystems, the complex life cycles of many NDF (aquatic lifestyles, obligate anaerobes, etc.) have led to dramatic undersampling and hindered their systematic study and adoption as research models^3,4^. However, many NDF lineages have retained features distinct from Dikarya, such as uneven rates of protist gene losses and the punctuated innovation of fungal-specific features in gene repertoires^17^. For instance, DNA and histone modifications display distinct conservation and functional roles. Repressive 5mC populates Dikarya genomes, whereas a DNA modification typically associated with bacteria, activation-associated 6mA, is predominant in and exclusive to non-Dikarya genomes^18,19^. Similarly, NDF show unique patterns of genome compaction and expansion, with genome sizes spanning over two orders of magnitude^20^. Resolving the dynamics of genome architecture and organization of information in NDF, epigenetic reprogramming and maintenance of genome integrity would fill a crucial gap in our understanding of lineage evolution since the last universal fungal ancestor.

Transposable elements (TEs) have been among the principal drivers of genome architecture and adaptation in fungi. Gene regulatory and defense mechanisms against TEs play highly diverse roles across the fungal kingdom. TEs have repeatedly escaped genome defense systems and are the major driver of fungal genome size expansion^21,22^. Moreover, TE dynamics have been shown to underpin adaptation to stress conditions, including thermal adaptation, antifungal drug exposure, and host infection in clinically and economically relevant Dikarya species^23–26^. Beyond recent transpositional activity, TE-derived co-option has been shown to play a pivotal role in the evolution of genic and structural functions, such as the emergence of point centromeres in yeast^27^. Furthermore, the characterization of fungal genomes has enabled groundbreaking discoveries, such as the description of giant transposons known as Starships in Pezizomycotina fungi, capable of transferring multiple genes between species and reaching sizes of nearly a megabase^28–30^. Recent community efforts have significantly expanded genome resources for non-Dikarya fungi^31^, providing an opportunity to leverage such datasets to recapitulate fungal genome dynamics beyond Dikarya.

Here, we analyze an extensive set of 163 genomes of NDF, along with 48 genomes covering non-fungal Holomycota, protists and Metazoa representatives, to retrace genome dynamics in the fungal kingdom since the Opisthokont ancestor. We assess which factors drove genome diversification and specific domain expansions. We reveal a multi-layered epigenetic landscape where TEs are tightly regulated by histone modifications and RNA interference. Furthermore, we uncover frequent horizontal transposon transfers across deeply divergent lineages, including cross-kingdom exchanges with protists. Our work establishes a comprehensive view of mechanisms underpinning genome architecture evolution across all major lineages of NDF and how mobile DNA impacted the early diversification of the fungal kingdom.

## RESULTS

### Highly diverse non-Dikarya genome architectures

To assess the extent of genomic variation across non-Dikarya fungal lineages in a broad phylogenetic context, we performed a large-scale comparative genomic analysis spanning 238 genomes across 13 fungal phyla/subphyla, including all publicly available non-Dikarya fungal genomes from each genus, 4 non-fungal Holomycota clades (Nucleariida (NU), Cryptomycota (CR), Microsporidia (MI) and Aphelidomycota (AP)) and 20 unicellular and multicellular Holozoa and Amoebozoa representatives (HO+AB) (**Figure 1a, Supplementary File 1**). Phylogenetic reconstruction was achieved with a set of 682 curated single-copy genes accounting for parsimony-informative sites and relative composition variability (see Methods), resulting in a topology consistent with recent reconstructions^15^. Genome sizes of non-Dikarya fungi and related non-fungal Holomycota range from 2 megabases (Mb) to over 700 Mb in some parasitic Entomophthoromycotina (EN) species and plant-associated Glomeromycotina fungi (GL) and Mucoromycota (MU) (Endogonales), comparable to genomes of complex metazoans. These expansions are driven by a remarkable abundance of TEs in their genomes (**Figure 1a**). All fungal and non-fungal Holomycota carry a high proportion (range 16-62 %, median 29.6 %) of genes that lack any annotation of conserved domains, which contrasts with well-studied animal models (**Figure 1a**).

**Figure 1.**
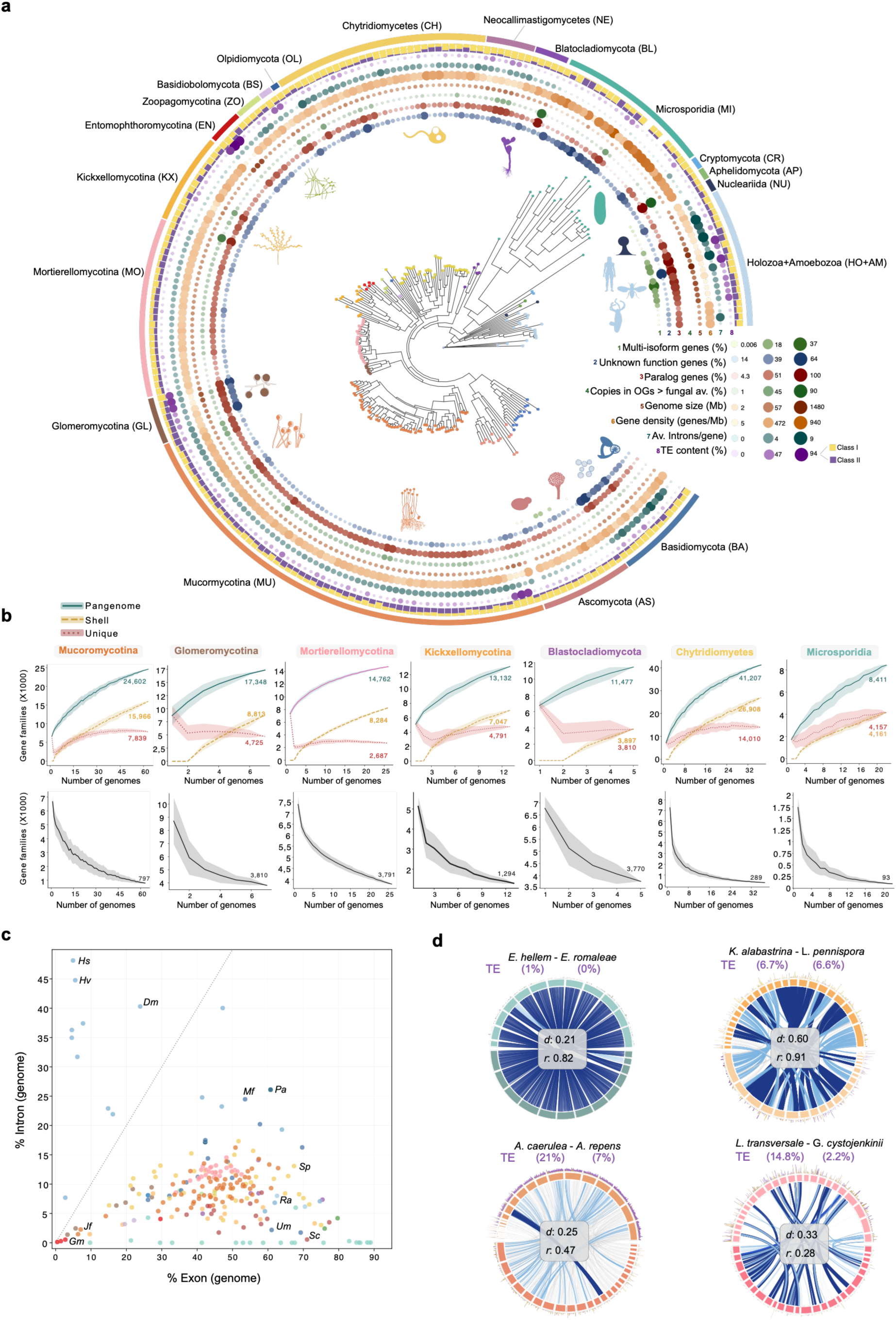
Genome-wide comparative genomics of NDF and related eukaryotes. (a) Maximum-likelihood phylogenetic tree of all 238 genomes. Tips are colored by clade and the abbreviations are labeled on the outer ring. (b) Pan-genome accumulation curves. In the top panels, each line shows the mean (± 1 s.d.) number of orthogroups classified as pangenome (≥ 1 genome), core (all genomes), shell (≥ 2 but < n genomes), or unique (1 genome) as a function of genome count, shown per major taxonomic group. Below, the core pangenome decay curve for each clade is indicated. (c) Scatter plot of exon content (% exonic, x-axis) vs. intron content (% intronic, y-axis) per genome. Each dot represents one species; color indicates phylogenetic clade. Note that TE-dominated genomes such as EN and Endogonales (MU) display both very reduced intron and exon content. Species abbreviations: Hs (*Homo sapiens*), Hv (*Hydra vulgaris*), Dm (*Drosophila melanogaster*), Mf (*Mrakia frigida*), Pa (*Parvularia atlantica*), Gm (*Gigaspora margarita*), Jf (*Jimgerdemannia flammicorona*), Sp (*Spizellomyces punctatus*), Um (*Ustilago maydis*), Sc (*Saccharomyces cerevisiae*), Ra (*Rozella allomycis*). (d) Circos microsynteny diagrams for four selected species pairs. Ribbons link collinear ortholog pairs; ribbon color corresponds to the scaffold in the left (reference) species. Corrected syntenic ratio (*r*) and patristic distance (*d*) are indicated for each pair. Total genomic TE content (%) is indicated for each species.

Dikarya fungal clades also display a characteristically high proportion of duplicated genes, associated with segmental and whole-genome duplications^32–34^. Comparative analysis of gene families in NDF clades further revealed that certain Blastocladiomycota (BL), Kickxellomycotina (KX) and MU species have undergone a substantial copy number expansion for a large proportion of their gene families (BL = 24 %, KX = 19.7 % and MU = 25.2% of families). Beyond gene family expansions, greater organismal complexity was also associated with gene family diversity. We conducted a pan-genome analysis of orthologous families for non-Dikarya clades and found the highest gene family diversity in MU and chytrids (CH), with up to 41,000 different gene families and only 289 core genes (*n=*37 genomes; **Figure 1b**). A very low number of core genes was also detected in MI species (*n*=22), indicating that the clade comprises both highly reduced and divergent genomes (**Figure 1b**). Despite the reduced sampling compared to Dikarya species, these results highlight substantial intraphylum diversity. Furthermore, this diversity is far from being assessed to saturation, and diversity metrics will likely increase considerably as more genomes become available. A further correlate of organismal complexity is higher rates of alternative splicing and gene intronization^35^. Although multi-isoform data are sparse for fungi, intron/exon gene structures are highly diverse, with some chytrids (CH), MU and Basidiomycetes (BA) (Dikarya) exhibiting animal-like high intron densities, while others are entirely devoid of introns, such as some Microsporidia (MI) genomes (**Figure 1a, 1c and Supplementary Figure 1a**).

An important feature of genome evolution is the retention of synteny blocks. We analyzed synteny conservation using the collinearity of orthologous gene sets. Since synteny is expected to decay as a function of divergence time, we calculated synteny conservation ratios (*r*) normalized by phylogenetic distance (*d*). Compact, repetitive element-free MI genomes displayed a high degree of synteny (**Figure 1d**), whereas pairs with comparable phylogenetic distances but higher TE content displayed a greater degree of syntenic block disruption (**Figure 1d**). Remarkably, within-phylum synteny turnover was particularly pronounced in clades with highly variable TE content, such as Neocallimastigomycotina (NE), Zoopagomycotina (ZO) and Mortierellomycotina (MO) (**Supplementary Figure 1b and 1c**). Collectively, these findings highlight the remarkable genomic heterogeneity within and between NDF lineages as well as the differentiation from their non-fungal relatives. Genome dynamics markedly surpass what was known for Dikarya lineages and implicate mobile elements as a major source of lineage diversification within the fungal kingdom.

### Idiosyncratic TE dynamics among non-Dikarya lineages

Given the apparent impacts of TEs on genome diversification in NDF, we next generated a high-quality annotation of these mobile elements for each species using a curated, kingdom-wide library of TEs^36^. Annotations were further refined using *de novo* annotation and curation of elements from non-fungal species included in this work (see Methods). Consistent with observations in other lineages, genome size positively correlated with overall TE content (Pearson’s *r* = 0.72), with LTR elements showing the tightest association (**Supplementary Figures 2a and 2b**). Notably, total TE content also positively correlated with intron density in non-Dikarya fungi, with LTR elements again displaying the strongest positive correlation (Pearson’s *r* = 0.19) (**Supplementary Figure 2b**). This underscores that TEs are likely the most significant determinants of genome size variation in non-Dikarya Holomycota lineages. Genomes of unicellular protists, other Metazoa, as well as other NDF representatives, exhibited the most extreme variation in TE abundance (**Figure 2a and Supplementary Figure 2a**). Within specific clades, distinct outliers with higher TE content emerged, such as the Endogonales in MU, *Lunasporangiospora selenospora* in MO, and *Smittium mucronatum* in KX (Figure **2a** **and Supplementary Figure 3a**). At the compositional level, non-Dikarya Holomycota displayed striking diversity both between and within clades (**Supplementary Figure 3b**). While LTR elements populated the TE-dense genomes of EN and NE, LINE elements occupied a significant proportion of the TE landscapes in MU and KX. Similarly, Helitrons (rolling-circle transposons) were prominent in several MU and MI genomes (**Figure 2a and Supplementary Figure 3a**). Remarkably, while some clades were dominated by a narrow set of TEs, MO and MU, followed by CH, showed the highest TE diversity among all studied groups, with several NDF representatives accumulating a larger number of different TE families than representative Dikarya lineages and comparable to Metazoa (Shannon H′ metrics; **Figure 2a and Supplementary Figure 3c**).

**Figure 2.**
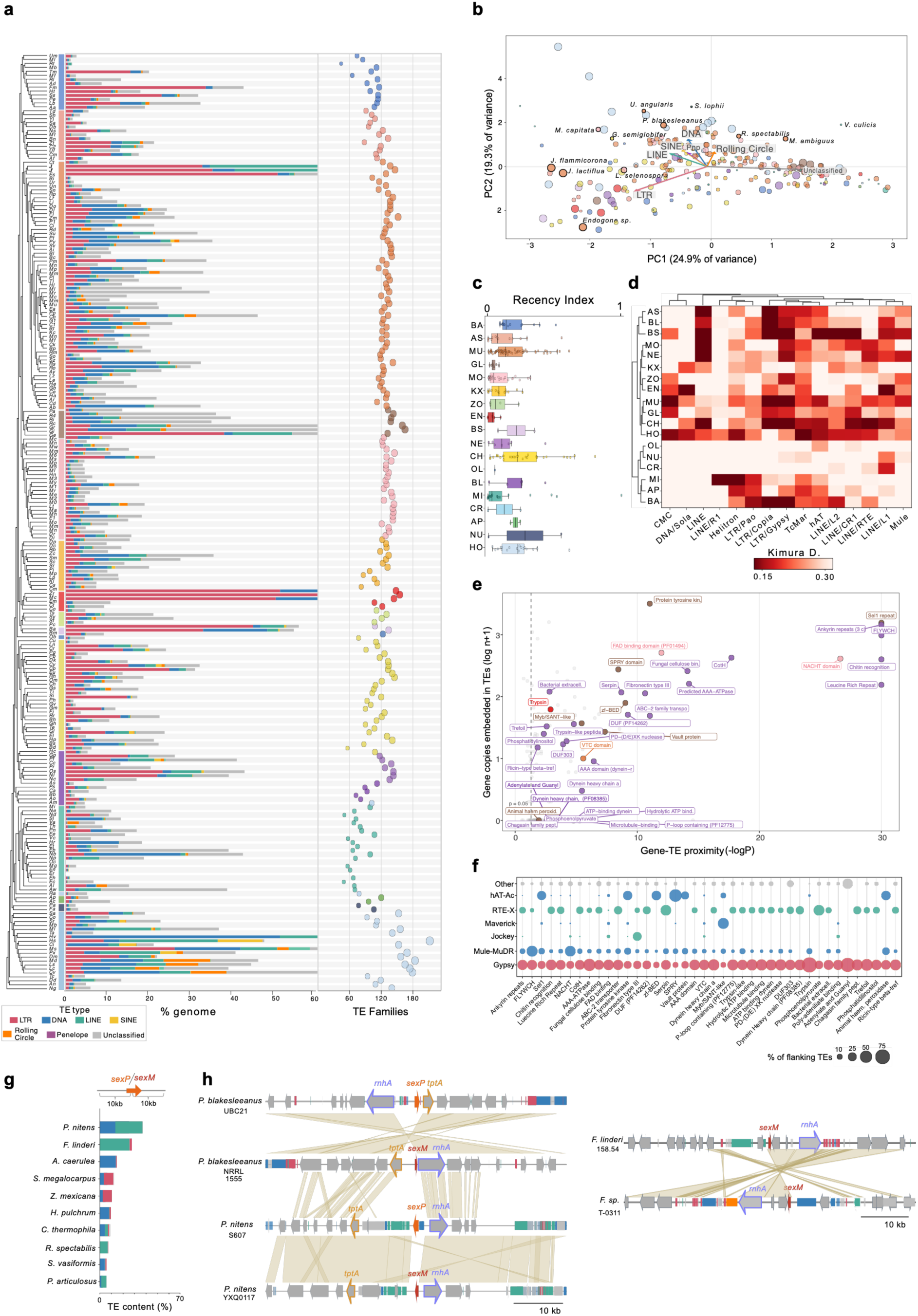
Role of TEs in genome diversification. (a) Cladogram of 238 species with TE class composition and the total number of different TE families for each species. (b) PCA of TE class composition across 238 species. Each dot represents one species colored by clade. Outlier species in their clade composition are labeled. Dot size is proportional to genome size. (c) Boxplot of the Recency Index (RI) per clade. Each box shows the interquartile range; whiskers: 1.5 × IQR; outliers: individual points. (d) Heatmap of Kimura divergence summary of TEs across 238 species, showing the mean Kimura divergence per TE superfamily. Species and superfamilies are ordered by Euclidean hierarchical clustering. (e) Expanded domains ordered by their gene–TE proximity significance (−log₁₀ BH-corrected Mann–Whitney *p*, expanded-clade gene copies vs. a size-matched random background in the same genome) and embedment within a TE annotation. (f) TE family frequency in symmetric flanks for the 41 domains in panel e. Bubble size shows % of each particular family in the flanking regions of genes containing expanded domains. (g) Total TE content (% occupied of the 20 kb flanking region) and class proportion in MU sex loci. (h) Genomic architecture of the sex locus in *Phycomyces* and *Fennellomyces*, showing the gene content and orientation of the mating-type region and TE distribution.

Non-Dikarya are sister lineages to Dikarya fungi emerging from an Opisthokont common ancestor shared with animals. While these evolutionary relationships are reflected at the gene family level^17^, a principal component analysis (PCA) of TE class composition revealed a complex, mixed landscape without clear grouping by subkingdom or clade, demonstrating that TEs diversified idiosyncratically in NDF (**Figure 2b**). Species with LTR TE expansions showed concomitant genome expansions, such as the insect-associated EN, NE and MU, whereas genome expansion in GL was mostly driven by TEs of unknown categories. In non-fungal clades, DNA elements drove most genome expansions (**Figure 2b**). Moreover, we identified TE composition outliers in diverse clades, such as LINE expansions in *Jimgerdemannia flammicorona* and *J. lactiflua*, SINE expansions in *Phycomyces blakesleeanus* and *Mortierella capitata*, and a pronounced expansion of Helitrons in *Radiomyces spectabilis*, *Mucor ambiguus*, and the microsporidian *Vavraia culicis* (**Figure 2b and Supplementary Figure 3b**), where these elements underpinned the most recent TE bursts (**Supplementary File 2**). TE activity leaves specific footprints in genomes depending on the epoch of activity^37^. Signatures of very recent or ongoing TE bursts were found in the human pathogen *Rhizopus delemar*, several CH species, *V. culicis*, and the microsporidian *Enterocytozoon bieneusi* (**Supplementary File 2**). Beyond these species, MU, Basidiobolomycota (BS), CH, and BL also exhibited high rates of recent TE insertions. Conversely, groups with highly TE-rich genomes (NE, EN, GL) showed lower rates of recent insertions, suggesting that these genome expansions were successfully counteracted by genomic defenses (**Figure 2c and Supplementary File 2**). In Dikarya genomes, most recent activity stemmed from LTR elements. In contrast, CMC elements caused the most recent TE activity in MU; R1 (LINE) and Helitrons dominated in MI; L1 (LINE) led activity in the non-fungal Holomycota clades NU and CR; and Sola (DNA transposons) and L1 elements dominated the recent TE activity in NE (**Figure 2d**).

Beyond characterizing the compositional landscape, we analyzed how TE burst dynamics contributed to genome evolution and lineage diversification. Following the discovery of high degrees of paralogy and gene family expansions, we performed a systematic analysis of expanded protein domains in non-Dikarya fungi at the species and clade level (**Supplementary File 3**). We focused on non-TE protein domains encoded within TEs, TEs positioned between tandem duplicates, and gene-TE proximity enrichment. We found evidence that 41 out of 255 expanded protein domains were associated with TE activity. These included the trypsin domain in EN, tyrosine kinase in GL, and the spore coat CotH domain and the flagella-related protein dynein, among others, in NE (**Figure 2e, Supplementary File 3, and Supplementary Figures 4a and 4b**). Domain expansions were frequently associated with Gypsy (LTR), the FLYWCH domain with MULE-MuDR TEs, the SPRY domain with hAT-Ac, and the Myb/SANT domain in GL with Maverick DNA transposons (**Figure 2f**). Together, these results demonstrate that TE activity and genome expansion dynamics are intimately linked to clade- and species-specific protein domain expansions defining the functional repertoire of non-Dikarya Holomycota lineages.

Genes controlling sex or mating type remain largely uncharacterized in NDF, with the exception of the HMG-box protein (sexP or sexM) and flanking genes in MU^38^. Given that the mating type locus can be structurally diverse and repetitive, we comprehensively analyzed these regions for TE content (**Figure 2g and Supplementary Figure 4c**). In addition to the inversion between the opposite mating types of *P. blakesleeanus* NRRL 1555 and UBC21, we detected TE remnants between *sexM* and the helicase gene *rnhA* in UBC21. Moreover, the sister species *P. nitens* displayed a full-length LINE inserted between the *sexM*/*sexP* locus and the triose phosphate transporter gene *tptA* (**Figure 2h**). *Fennellomyces linderi* exhibited substantially higher polymorphism at the mating type locus compared to its sister species, consistent with the recent transposon burst observed in *F. linderi* (**Supplementary File 2**). The TE-dominated mating-type locus contains LINE and LTR elements in one genotype, and a Helitron insertion alongside DNA transposons in another (**Figure 2h**). Overall, these results show that TE insertion activity is a recurrent source of structural variation at the mating-type locus beyond Dikarya fungi.

### Multi-layered epigenetic mechanisms regulate TE activity

Epigenetic mechanisms have coevolved with TE activity as key genomic defenses^39^. We focused on MU as the best-represented non-Dikarya clade to investigate how TE activity is dynamically regulated across developmental stages and in response to environmental conditions. First, we analyzed the repression-associated DNA modification 5-methylcytosine (5mC). Both DNMT1 and DIM2, the core components responsible for 5mC deposition, have undergone recurrent, parallel losses in MU, leading to few species retaining both components (**Figure 3a and Supplementary File 4**). Among them are the Endogonales and *P. blakesleeanus*, the latter being the only species where high CpG methylation levels have been experimentally validated^18,19^. Interestingly, species retaining the methylation machinery exhibited higher TE contents than those lacking one or both components (**Supplementary Figure 5a**). This is consistent with this machinery being important in TE-rich genomes and having potentially co-evolved with TE activation dynamics. Although 5mC has not been experimentally verified in Endogonales, they displayed a drastically reduced GC content of TEs. Hence, methylation has likely contributed to the inactivation and domestication of TE sequences via the deamination of methylated cytosines (**Figure 3a**). This was also supported in *P. blakesleeanus*, where younger TEs were enriched in 5mC (**Figure 3b and Supplementary Figure 5b**).

**Figure 3.**
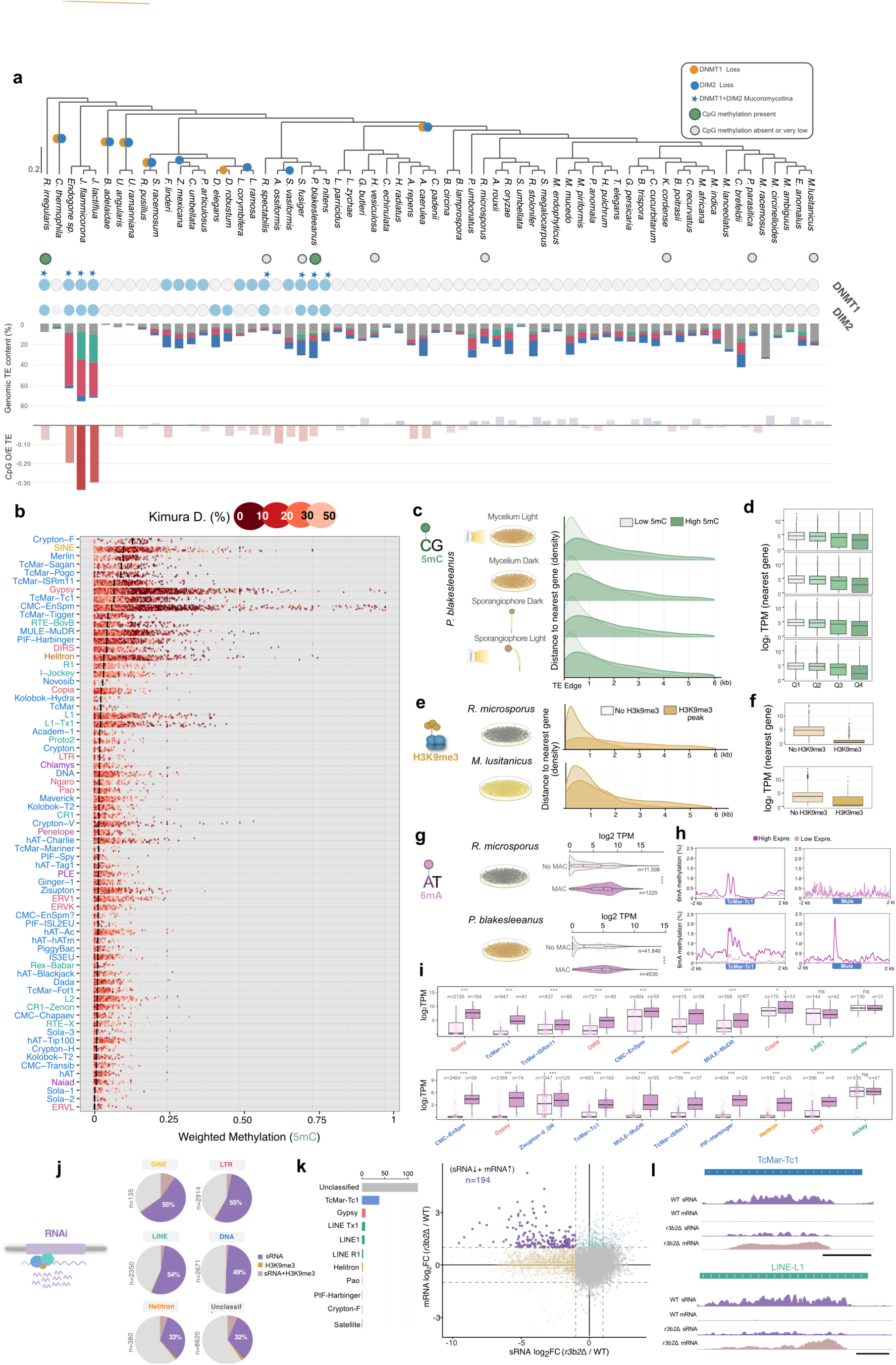
Epigenetic regulation of TEs in Mucoromycota. (a) Maximum-likelihood phylogenetic tree of MU species (rooted to *R. irregularis*) showing conservation of DNMT1 and DIM2 for each species. Losses of the 5mC methylation machinery are indicated as yellow (DNMT1) and blue (DIM2) circles. Experimental evidence of the presence of CG methylation is indicated for each species (green = present, light green = absent, no dot = no experimental data). TE content and composition, as well as the Observed/Expected ratio of CG content of TE copies (considering per-species GC content), are also indicated. (b) 5-methylcytosine (5mC) per-copy weighted mean by TE family and Kimura divergence age class. Only TE families with ≥ 50 covered copies were considered. (c) Cumulative distribution of gene distance from the nearest full-length TE stratified by per-copy 5mC methylation quartile (Low/High). Kolmogorov–Smirnov tests between High and Low 5mC (p <0.01 for the four growth conditions). (d) Expression of genes adjacent to TEs, stratified by methylation quartile as described before. Expression was computed as log2TPM. Groups were compared by Benjamini–Hochberg-corrected Mann–Whitney U tests (p <0.01 for the four growth conditions). (e) Cumulative distribution of gene distance from the nearest full-length TE stratified by per-copy H3K9me3 methylation quartile (Low/High). Kolmogorov–Smirnov tests between High and Low occupancy (p <0.01 for the two species). (f) Adjacent gene expression by H3K9me3 peak presence/absence. Statistical comparisons were performed as in d (p <0.01 for the two species). (g) Violin plot comparing expression of TEs with ≥ 1 MAC in the TSS window (−150 to +400 bp) versus TEs without MACs, for *P. blakesleeanus* and *R. microsporus* grown under mycelium dark conditions. (p < 0.001 for both species, Mann–Whitney U test (two-sided) BH-corrected). (h) 6mA occupancy profile of TcMar-Tc1 and MULE-MuDR TEs. Each line represents one expression quartile. (i) Expression (log2TPM) of TE copies grouped by the presence/absence of MAC at the TSS (−150 to +400 bp). (Wilcoxon BH-adjusted * p<0.05 ** p<0.01 *** p<0.001, ns = not significant). (j) Pie charts showing the proportion of TEs targeted by RNAi, H3K9me3 or both in *M. lusitanicus*. (k) Scatter plot of sRNA abundance (RPM, y-axis) vs. TE copy expression (RPKM, x-axis) in WT vs *r3b2* mutant strain of *M. lusitanicus.* Top left corner groups TEs (194) with reduced sRNA accumulation and increased mRNA accumulation in the mutant. Classification of the candidate target TEs is indicated in the bar chart. (l) Genome snapshots of two RNAi target candidate TEs, including sRNA and mRNA tracks for WT and *r3b2* mutant strain. Scale bar = 500 bp.

**Figure 4.**
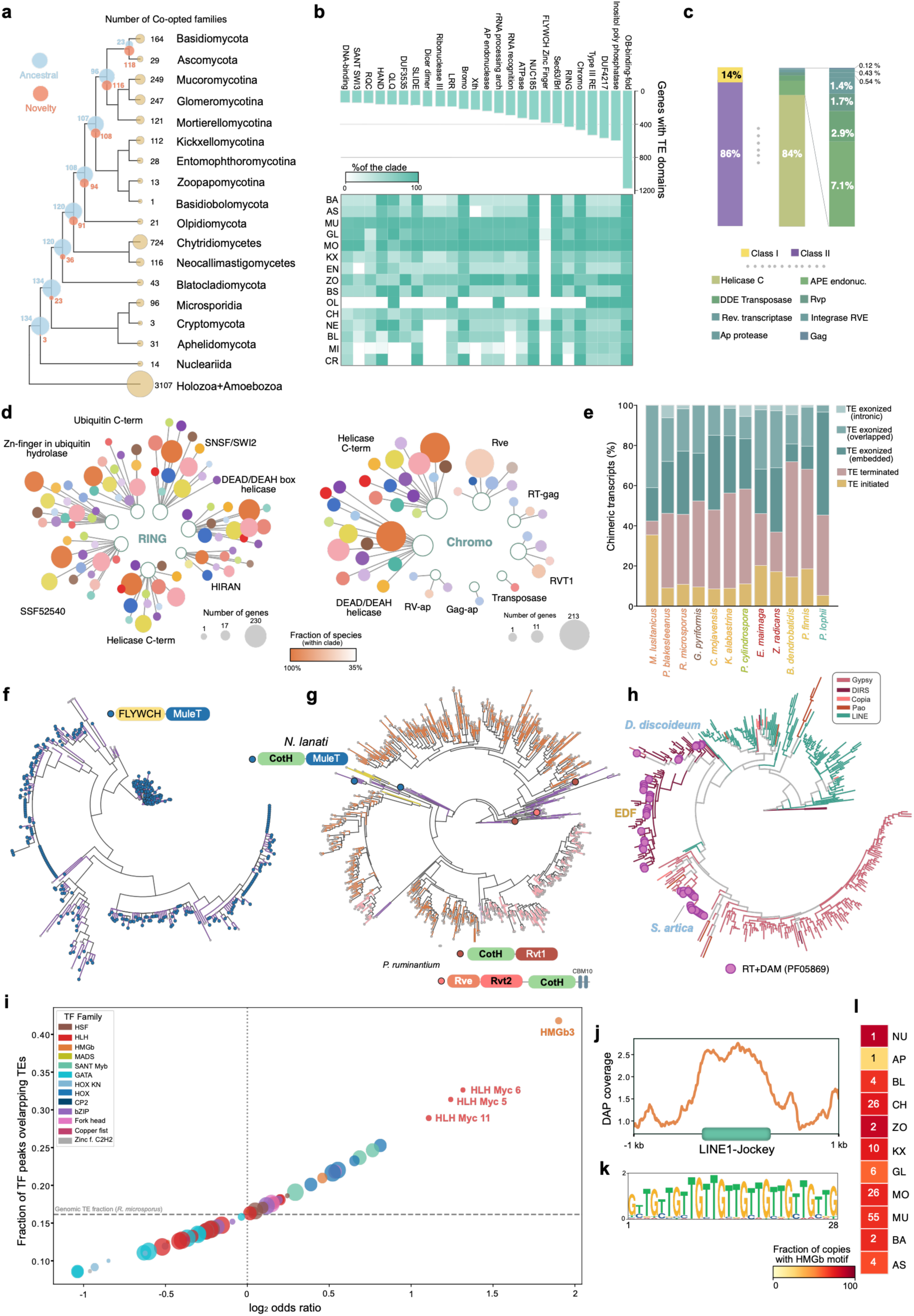
Functional and structural TE co-option across NDF. (a) Gains of new and losses of ancestral (shared with HO+AB) co-opted genes. At each ancestral node, the number of co-option orthogroup families conserved from the ancestor and the number of newly acquired families are annotated. At the tip of each branch, the total number of distinct orthogroups with co-opted TE-derived genes within that clade is indicated. (b) Most prevalent non-TE Pfam domains in co-opted genes per clade (top) and heatmap indicating the proportion of species in each clade in which this co-option is detected (bottom). (c) Stacked bars show the fraction of TE class and TE domains involved in co-opted genes. (d) Fusion protein co-option network. Each node shows one interacting domain with RING (left) and Chromo (right) domains. Edge width for each domain shows the number of species with this fusion, and color transparency indicates the fraction of the species for each clade with the indicated domain co-occurrence. (e) Stacked bar chart showing the proportion of different types of chimeric junctions for all TE-containing transcripts. (f) Phylogenetic trees of FLYWCH domain-containing proteins, restricted to Neocallimastigomycota. (g) Phylogenetic trees of fungal CotH proteins across clades. (h) Phylogenetic tree of Reverse Transcriptase (RT) domains. Pink circles indicate RT domains in frame with a DAM domain. (i) Bubble chart of peak enrichment of TF binding using DAP-seq data. Bubble size indicates the number of TF–TE combinations significant after BH correction. (j) Occupancy profile of HMGb (DAP IP/Input control) over LINE/I-Jockey copies in *R. microsporus*. (k) MEME motif logo for the HMGb3 binding motif. (l) Heatmap indicating the fraction of LINE/I-Jockey copies in each clade containing the HMGb3-recognized motif.

Given that chromatin modifications can leak into flanking regions, we examined whether TE-associated epigenetic marks influenced the transcriptional activity of neighboring genes. We analyzed a comprehensive dataset of both *R. microsporus* and *M. lusitanicus* covering developmental (mycelium/sporangiophore) and environmental (dark/light) conditions. Genes adjacent to densely methylated TEs showed lower expression than those near poorly or unmethylated TEs (**Figures 3c and 3d**). Gene silencing adjacent to TEs was marked by the repressive histone modification H3K9me3 (**Figures 3e and 3f**). Notably, despite performing similar repressive roles, H3K9me3 preferentially targeted TcMar TEs, whereas 5mC was specifically enriched over LINE elements in *P. blakesleeanus* (**Supplementary Figure 5d**). Then, we investigated whether 6-methyladenine (6mA), generally associated with active gene expression^19,40,41^, was associated with transcriptionally active TEs. Notably, TEs harboring Methylated Adenine Clusters (MACs) were more transcriptionally active than those lacking them, and this effect was most pronounced for DNA transposons (**Figure 3d and 3g**). Highly expressed TEs exhibited characteristic 6mA enrichment flanking the transcription start site (TSS), mirroring non-TE genes (**Figure 3h**). We found that 6mA accumulation and expression of CMC-EnSpm (*r* = 0.38, *p* < 0.004), Helitron (*r* = 0.47, *p* = 0.0002) and TcMar-Tc1 (*r* = 0.47, *p* < 0.001) families were positively correlated in *P. blakesleeanus* but not in *R. microsporus* (Fisher z-transformation; **Supplementary Figure 6**). This highlights a potential species-specific role for 6mA in stimulating TE transcription. Conversely, increased 5mC levels were associated with lower expression of TcMar, RTE-BovB, and Gypsy copies in *P. blakesleeanus* (**Supplementary Figure 6**). Pearson *r* = 0.39, one-tailed *p* = 0.004, n = 45 copies; and *r* = 0.58, *p* = 0.003, n = 21 copies, respectively; 95 % CI by Fisher z-transformation Since epigenetic marks show preferential deposition over specific TE families, we examined whether this impacts the chromatin architecture of TE-rich regions. In vivo MNase-seq-generated data from MU and Dikarya fungi show that nucleosomes in TE-rich regions form tighter arrays with higher spacing variability compared to genic sequences with more regularly spaced and dispersed nucleosome organizations (**Supplementary Figure 7a)**. Conversely, sequence-based predictions of nucleosome occupancy with models trained on fungal genomes^42^ failed to replicate this distinct pattern (**Supplementary Figure 7b and 7c**), suggesting that epigenetic modifications associated with TE-rich regions, rather than the nucleotide sequence of TEs, direct nucleosome architecture in these genomes. Since 6mA has been linked to nucleosome positioning in eukaryotes^40^, we compared nucleosome profiles over MAC-bearing and unmethylated TEs. MAC-marked copies displayed more defined and consistently positioned nucleosome arrays, which favor RNA polymerase accessibility and likely explain their elevated transcriptional activity (**Supplementary Figure 7d and 7e**).

Beyond investigating transcriptional control of TE-rich regions in MU, we assessed post-transcriptional regulation mediated by RNAi. Integrating our curated TE annotations with small RNA (sRNA) sequencing data revealed that a large proportion of TEs accumulate targeting sRNAs in *M. lusitanicus*, with most H3K9me3-marked TEs being co-targeted by sRNAs (**Figure 3j and Supplementary Figures 5e and 5f**), which suggests a tight coordination between chromatin-level repression and RNAi-mediated surveillance over the same TE loci. Furthermore, analyzing a mutant of the non-canonical RNAi pathway, *r3b2*Δ^43^, we identified 194 putative TE targets that exhibited significantly increased mRNA accumulation coupled with depleted sRNA levels (**Figures 3k and 3l**). This supports the idea that RNAi-mediated post-transcriptional control is an important factor in TE activity. Collectively, we unravel an epigenetic TE regulatory system in MU that is multilayered and differentially targets classes of TEs. While 5mC, H3K9me3, and RNAi converge on TE silencing, 6mA acts as a lineage-specific activation counterforce. Such diversity of control mechanisms mirrors the compositional complexity of EDF mobilomes. Our insights underscore the distinct co-evolutionary arms races of TEs and host genomes playing out across deeply divergent fungal lineages.

**Figure 5.**
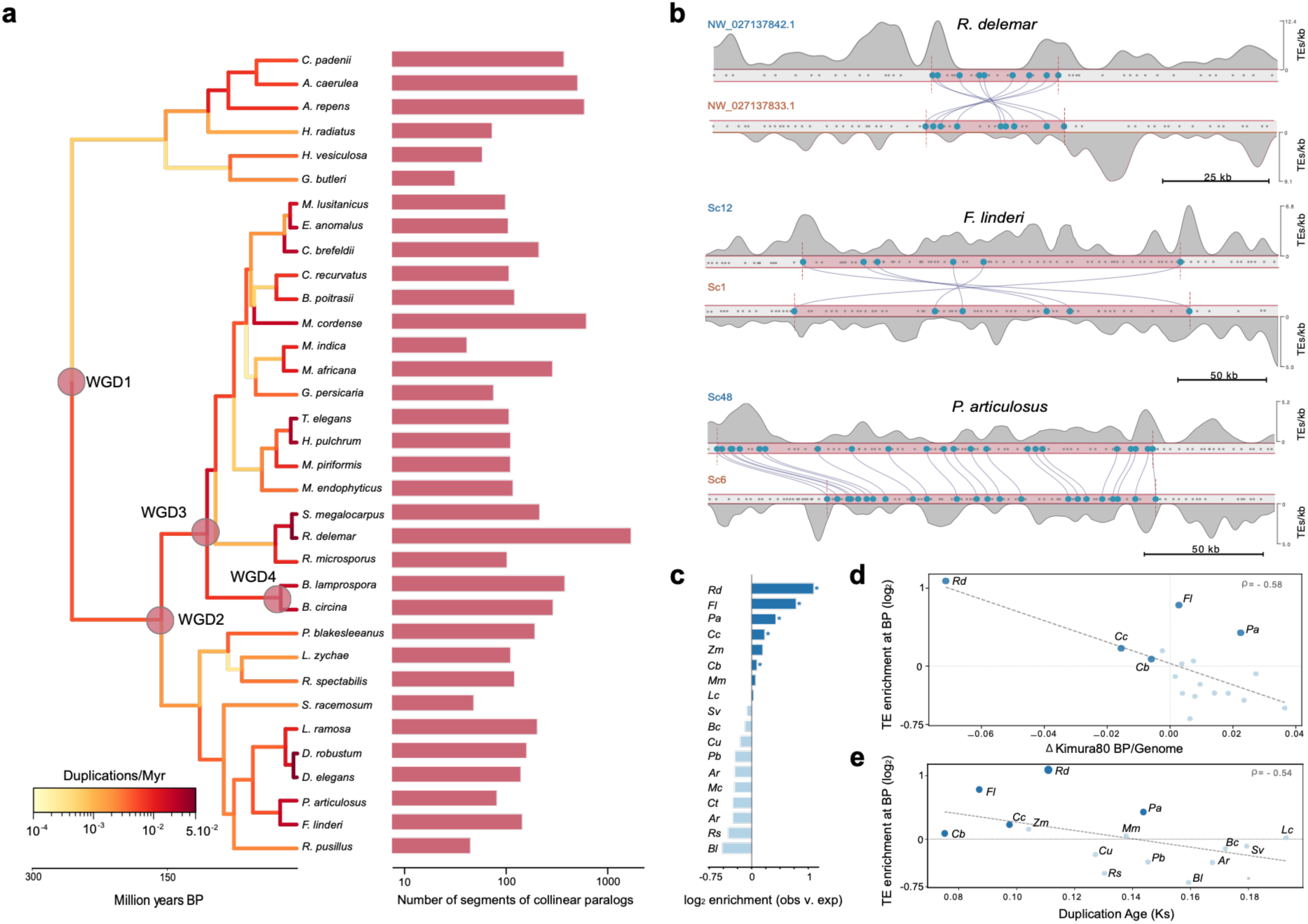
WGD and TE dynamics in Mucoromycota. (a) MU phylogenetic tree (limited to species with < 200 scaffolds). Normalized duplication rates per phylogenetic node were computed as the number of duplication events mapped to each node. Tree branches are colored on a continuous heatmap with normalized duplication rate, giving a visual proxy for duplication events per million years from the dated tree generated by Szánthó et al.16. Total count of multiplicons for each species is indicated. (b) Representative loci showing multiplicon collinearity and TE density tracks. (c) TE enrichment at multiplicon breakpoints and flanking regions (10 kb) calculated as log2FC over randomized windows. Asterisks indicate species with significant TE enrichment in these regions. (d) ΔKimura80 (mean Kimura divergence at breakpoints − genome-wide mean; Δ < 0 means younger insertions at breakpoints) vs TE enrichment at breakpoints (log₂FC over randomized windows). (Spearman ρ = −0.58, p = 0.012). (e) Spearman correlation between anchor-pair Ks and TE superfamily burst mode, for species with reliable Ks estimates vs TE enrichment at breakpoints (log2FC over randomized windows). (Spearman ρ = −0.54, p = 0.028).

### Recurrent co-option of mobile elements drives functional and structural innovation in NDF genomes

Given the widespread association between TE activity and clade- and species-specific domain expansions across NDF genomes, we next investigated the broader functional implications of these mobile elements through a systematic analysis of TE-driven gene co-option (**Supplementary File 5**). Gene families co-opted by TEs prior to the split of Holozoa and Amoebozoa have been progressively lost in the Holomycota lineage. However, we find that the loss has been heterogeneous, with NDF clades retaining a large fraction of these ancestral co-opted families and only 23 persisting in Dikarya (**Figure 4a and Supplementary Figure 8a**). Our findings are in contrast with previous screens for gene co-option in the fungal kingdom, which undersampled NDF lineages (less than 6% of the analyzed organisms)^44^. Most notably, we find an expansion of the FLYWCH domain and several domains of unknown function (DUFs) that are rare or absent in Dikarya (**Figures 4b and 4c**). Co-option events were dominated by Helicase C domains, followed by endonuclease and DDE transposase domains (**Figure 4c**). Interestingly, while co-opted genes in Holozoa and Amoebozoa (HO+AB) contain fewer introns than regular genes, a feature shared with Dikarya (BA), most co-opted genes in NDF are remarkably intron-rich (**Supplementary Figure 8b**). Examination of co-occurrence networks for the most frequently co-opted domains revealed clade-specific architectures; for instance, the Chromo domain is frequently fused to Rve domains in BA, but couples with transposases in EN or Rvt1 in MU (**Figure 4d)**. Co-opted DUFs formed complex co-occurrence networks. DUF4217 was particularly diversified, with TE domain fusions as well as kinesin or cold-shock DNA-binding domain fusions (**Supplementary Figure 8c**). Transcript-level analysis of chimeric RNAs revealed a predominance of TE-terminated transcripts across lineages, except in *M. lusitanicus* (MU), where nearly 40% of chimeric transcripts were initiated by TEs (**Figure 4e**).

We next assessed recurrent fusion events involving FLYWCH (a C2H2 zinc finger), CotH, and NACHT domains by searching for open reading frames (ORFs) encoding these focal domains in addition to a TE domain (**Supplementary Figure 8d and Supplementary Table 1**). The FLYWCH domain has extensively expanded in NE, as well as in some CH and GL species, frequently co-opting a MULE transposase (**Figure 4f and Supplementary Figure 9**). We propose that this phylogenetically widespread domain fusion should be identified as a novel subfamily of DNA transposons, termed Genomic Non-Dikarya MULE Elements (GNOMEs). Additionally, two copies of NACHT domains were co-opted within the same ORF alongside an LTR transposon Rvt1 domain in *Dissophora decumbens* (MO) and *L. selenospora* (MO) (**Supplementary Figure 8e and Supplementary Table 1**). The CotH domain, a critical component of spore coat proteins in MU that mediates fungal-mammalian host interactions during infection^45,46^ and whose copy number correlates with virulence^47^, showed complex co-option patterns, including in anaerobic gut fungi (NE). Similar to the FLYWCH co-option, CotH was co-opted by a MULE transposase in *Neocallimastix lanati* (**Figure 4g, Supplementary Table 1**). *Pecoramyces ruminantium* harbors two distinct co-options with either Rvt1 or Rve+Rvt2 (**Figure 4g**). Remarkably, the latter ORF also contained two carbohydrate-binding modules (CBM10) (**Figure 4g and Supplementary Table 1**). Functional implications of such chimeric architectures found in frequent inhabitants of ruminant microbiota remain unknown. Next, we investigated co-option of domains linked to DNA methylation (DNMT, MTA-70, N6-MTASE, and DAM). Consistent with recent findings in non-fungal eukaryotes^48^, we observed that bacterial DAM methyltransferase domains are frequently co-opted by transposons in NDF (**Figure 4h and Supplementary Figure 10a**). Expanding beyond the Gypsy and DIRS fusions found in *Sphaeroforma arctica* and *Dictyostelium discoideum*, we detected DIRS-mediated co-option of DAM across GL, KX, MO, MU, and ZO clades. This suggests that this transposon class may present unique competence for domain co-option in NDF but not in Dikarya fungi (**Figure 4h and Supplementary Figure 10a**).

Beyond driving gene function diversification, TE sequences can also be co-opted as *cis*-regulatory elements or transcription factor (TF) binding sites (exaptation). Such co-option can subsequently evolve into structural regions such as centromeres^27^. Leveraging a TF-binding dataset recently generated for the NDF *R. microsporus*^49^, we screened 58 different TFs for binding preferences across TE sequences. We detected a pronounced binding enrichment of three helix-loop-helix (HLH) TFs over Chlamyvir Gypsy elements (**Figure 4i and Supplementary Figure 10b**) and, even more strikingly, the binding of a high-mobility group (HMG) TF (HMGb3) onto LINE-Jockey elements (**Figures 4i and 4j**). The repetitive motif recognized by HMGb3 (**Figure 4k**) was widely conserved across other LINE-Jockey copies (**Figure 4l**). Furthermore, the motif was also found in all 44 copies of a class of centromeric TEs (GremLINEs) identified in *M. lusitanicus*^50^ (**Supplementary Figure 10c**), suggesting a conserved structural role for these mobile elements in genome architecture in NDF. Together, our broad screen for functional co-option revealed novel domain fusions, new TE family associations, and the recurrent use of transposon-derived sequences as transcription factor binding sites, indicating that NDF evolution was facilitated by a diverse and previously unrecognized repertoire of TE neofunctionalization.

### TE-driven repeated whole-genome duplications in Mucoromycotina

Whole-genome duplication (WGD) is one of the major drivers of large-scale genomic variation in fungi^32^, including NDF^51,52^. We therefore investigated the extent to which TE dynamics may drive WGD events across NDF lineages. WGD has only been reported in the Mucoralean species *R. delemar*, with TE enrichment near duplicated blocks^51^. We expanded the search for WGD to newly available, highly contiguous genomes and used a synteny tracking framework based on collinear duplicated gene segments (multiplicons)^53^. The fine-grained analysis of duplication events was performed on the 34 most contiguous MU genomes (**Figure 5a, Supplementary File 6**). This resolved four WGD events (**Figure 5a**), which add to the two WGD events previously proposed for the *Rhizopus* clade and the inferred MU ancestral WGD event^54^. We detected high terminal duplication rates in *R. delemar* consistent with these previous reports, as well as in the closely related species *Syzygites megalocarpus*. Other species, including *M. lusitanicus* and *P. blakesleeanus*, also displayed elevated duplication rates coinciding with ancestral WGD events near the root of the clade, extending and refining previous observations on a more limited number of species^33^. We then analyzed whether segment duplication breakpoints and their flanking regions (10 kb) were populated by TEs. We detected an enrichment of TEs at the breakpoints and flanking regions of 5 MU species, including *R. delemar* (**Figure 5b and 5c**). Interestingly, this enrichment was not associated with total TE content in the genome, suggesting that TE accumulation at duplication boundaries has specific drivers rather than being a consequence of genome-wide TE content (**Supplementary Figure 11a**). To assess the specificity of this TE accumulation related to WGD events, we examined the TE family composition at breakpoints (**Supplementary Figure 11b**). TcMar elements (in particular TcMar ISRm11) recurrently colonized duplication boundaries in independent multiplicons of *R. delemar* and *Chaetocladium brefeldii*, with a high proportion of collinear fragments displaying TcMar ISRm11 copies both in their flanking regions and within the duplicated segment pairs themselves (**Supplementary Figure 11c**). This specific accumulation could be explained either by high element activity predating the WGD event and the elements contributing to the duplications, or by an insertion preference of the elements for these regions following the WGDs. In other MU representatives, no particular family enrichment was detected, with the exception of Ginger DNA elements in flanking collinear fragments of *Choanephora cucurbitarum* (**Supplementary Figure 11b and 11c**). Nevertheless, TE accumulation in these regions was positively correlated with the presence of younger copies across MU lineages (**Figure 5d**). Finally, we investigated the temporal components of TE accumulation in the collinear segments and found that TE enrichment was strongest in the youngest segment duplications, as estimated by silent site divergence (K_S_) (**Figure 5e**). This would be consistent with both rapid TE deactivation in these regions following an initial burst and recent TE activity driving accumulation at breakpoints. Our results establish that TE enrichment at breakpoints is a recurrent feature of WGD in MU and illuminate the molecular dynamics of a phenomenon with nearly no parallel in Dikarya.

### Horizontal transfer of TEs with non-fungal Holomycota and distant protists

Given the pervasive impact of TEs on NDF genome evolution and diversification, we assessed the evolutionary origins of TEs beyond vertical inheritance. For this, we leveraged our curated TE annotation to systematically infer horizontal transposon transfer (HTT) events across the phylogenetic breadth of our dataset, including non-fungal clades (**Figure 6a and Supplementary File 7**). Following earlier large-scale investigations of fungal HTT, primarily focused on Dikarya, we first calibrated baseline evolutionary divergence between each pair of analyzed clades using silent site divergence (K_S_) of orthologous genes. Then, we screened carefully curated TE ORFs to identify extreme lower-bound divergence outliers compared to genome-wide estimates. Among all TEs with strong HTT evidence, LINE elements accounted for the majority of the interclade transfers (**Figure 6b and Supplementary Figure 12a and 12b**). This is in contrast with Dikarya species, in which DNA elements dominated recent HTT events^55^. Among NDF, MU accounted for a large portion of the detected transfers, particularly with the closely related subphylum MO, but also with the more distant CH and BS lineages (**Figure 6c**). Next, we focused on cross-kingdom transfers between EDF and non-fungal clades. Interestingly, we detected evidence for Helitron transfer events between a CH species and the non-fungal Holomycota *Fonticula alba* (NU) (**Figure 6a, 6c and Supplementary File 7**) as well as an LTR element transferred between CH and AP (**Figure 6a, 6c and Supplementary File 7**). Strikingly, HTT also occurred between the choanoflagellate protist *Monosiga brevicollis* (Choanoflagellatea, HO) and CH and MO species, which diverged over a billion years ago^16^ (**Figure 6a, 6c and Supplementary File 7**). The transfers included a LINE element sharing >80% identity between a *Linnemannia* ancestor and the protist (**Figure 6d**). Our findings of cross-kingdom HTT between EDF lineages and other distant eukaryotes highlight the complexity of TE content dynamics. Given the propensity of TEs to drive genome structure evolution in early-diverging fungal lineages, broader searches for HTT evidence and biological mechanisms of transfer will be crucial.

**Figure 6.**
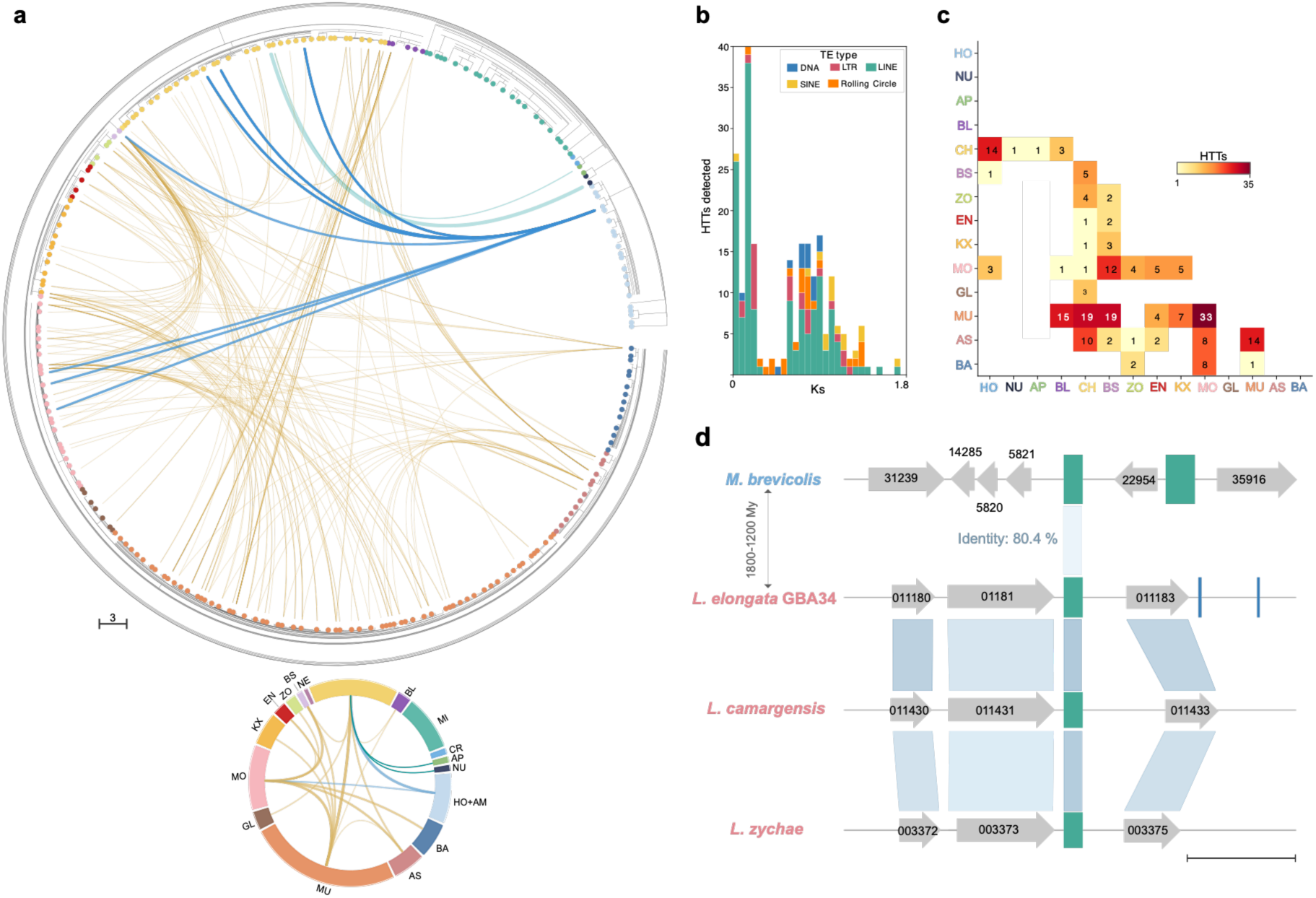
Evidence for widespread intra- and inter-kingdom HTTs. (a) 223 individual HTT candidates shown as links between species. Fungal transfers are indicated in yellow, fungal–non-fungal Holomycota in teal, and fungal–HO+AB transfers in blue. The 15 minimal transfer events accounting for vertical transmission are described in the diagram below. (b) Ks distribution of HTT candidates, stacked by TE class. (c) Heatmap of HTT candidate counts per clade pair. (d) Synteny analysis of *M. brevicollis (HO) and Linnemannia* species (MO) sharing a LINE TE. Scale bar = 1 kb.

## DISCUSSION

Early-diverging fungal branches and non-fungal Holomycota represent a critical repertoire of organisms for understanding major evolutionary transitions towards multicellularity and terrestrialization. Analysis of NDF lifestyles and their genome evolution illuminates these transitions independently of transitions in green and brown algae, as well as animals. Despite their importance and deep diversification, NDF have been largely neglected in favor of the ecologically more abundant Dikarya^20^. Here, we show that TEs followed highly varied trajectories in EDF genomes, even between closely related species.

We show that TE activity most likely had consequences for the evolution of specific lifestyles and protein domain expansions. Most notably, proteins such as trypsin in EN, Myb/SANT or protein tyrosine kinase in GL, and several distinct domains in CH and NE, such as dynein, CotH, or cellulose-binding domains, were co-opted or expanded during EDF evolution. Building on previous evidence^56^, we were able to directly associate the expansion of specific domains with TE activity. Gene co-option screens in fungi have typically underrepresented EDF^44^, as often <10% of total datasets stemmed from this group^20^. Our analysis provides an unbiased perspective on the evolution of domain architectures in EDF and contrasts these dynamics with patterns specific to Dikarya. In particular, we detected TE-associated domain expansions such as FLYWCH, occurring exclusively in CH, GL, and NE, or the specific CotH fusions detected in anaerobic gut fungi representatives (NE). Additionally, we found that domains involved in cellulose metabolism (CBM10) were encoded in the same gene as Rve, Rvt, and CotH. This finding opens an entirely new avenue to investigate pathogenesis-associated domains in Mucoromycota and their roles in cellulose processing in ruminant digestive tracts. The link to mobile element expansions underscores the highly dynamic nature of such domain architectures. Bacterial DAM adenine methyltransferase domains have been found to be co-opted by Gypsy and DIRS transposons in several eukaryotes^48^. Here, we show that this phenomenon is also widespread in fungi, with several NDF lineages carrying DAM. The absence of the domain in Dikarya reinforces the need to more systematically consider functional diversity in the fungal kingdom. Furthermore, the presence of DAM in some EDF clades is consistent with analyses of protein families revealing that EDF clades retained more ancestral features shared with protists than Dikarya fungi^17^.

Beyond their potential for gene family expansion and neofunctionalization, TEs can also provide specific binding sequences recognized by transcription factors^57^. Here, we leveraged the largest transcription factor binding dataset available for NDF and found that TEs are tightly associated with specific transcription factors. Our analyses revealed that the binding site of an HMGb3 transcription factor was frequently located in specific centromeric LINE elements of MU^50^, suggesting that TEs could play broad roles in structuring fungal centromeric regions. This is further supported by evidence of how yeast centromeres evolved from ancient TE sequences co-opted as centromeric protein-binding sites^27^.

Epigenetic mechanisms governing transcriptional and post-transcriptional activity of TEs represent a widespread phenomenon across eukaryotes^58^. Here, we found that conservation of the 5mC methylation machinery is directly associated with total TE content in MU. Loss of the machinery occurred in species with a lower abundance of mobile elements, whereas the machinery was strictly retained in TE-rich genomes. In addition to the transcriptional repression mediated by 5mC, we show that the constitutive heterochromatin mark H3K9me3 provides transcriptional repression in species lacking CG methylation. Beyond methylation, the RNA interference pathway serves as a widespread mechanism to target TE-derived mRNAs, especially those escaping H3K9me3 transcriptional repression. Both mechanisms showed strong convergence in TE targeting, which suggests that H3K9me3 serves as a guide for RNAi mechanisms, as has been reported for host genes^59^. Our analysis of 6mA, a DNA modification linked to transcriptional activation and found in some NDF but not in Dikarya^19,60^, revealed that it is associated with the transcription of specific DNA transposons. Strikingly, loci encoding the DAM domain, responsible for bacterial adenine methylation and co-opted by TEs in NDF, are densely covered by active 6mA marks in protists^48^. This is consistent with a model in which specific TEs evolved an escape mechanism to remain transcriptionally active in 6mA-competent genomes, thereby facilitating their expansion.

A major mechanism of eukaryotic lineage diversification is WGD. Despite a number of such events in Dikarya and NDF, WGDs have traditionally been challenging to detect and characterize^32^. A well-studied WGD occurred near the root of the Saccharomycota^61,62^ and recent studies expanded the catalogue to four additional lineages using novel approaches^53^. Further evidence pointed to WGDs in the Mucoralean species *R. delemar*^51^. Ancestral state reconstructions suggested at least one additional event in MU prior to this burst^33^. The reliance on only four MU genomes complicated these analyses considerably. Here, the increased availability of highly contiguous genome assemblies has enabled us to identify multiple ancestral WGDs throughout MU evolution. Moreover, the observation made in *R. delemar* regarding the accumulation of TEs at the boundaries of duplicated blocks was further confirmed here and extended to other MU species, providing robust evidence for recurrent WGDs in Mucorales. Our analyses revealed that TE dynamics follow temporal patterns, with the most recent blocks showing the most substantial colonization. This makes it possible to build mechanistic models of how peaks in TE activity may have favored the recurrence of WGDs in lineages.

Beyond functional consequences of TE activity in NDF genomes, we analyzed evidence for non-conventional inheritance through horizontal transfers. Based on a robust phylogenomic framework, we found an extensive set of transfer events, extending earlier work primarily focused on Dikarya fungi. We found LINE elements to be the most frequently transferred TE class between non-Dikarya clades. This stands in contrast to DNA elements dominating detected transfer events in a dataset comprising mostly Dikarya^55^. This highlights how both biological mechanisms of transfer and transfer competence of TEs themselves may substantially differ in EDF and be masked by analyses dominated by Dikarya genomes. Strikingly, our analyses revealed evidence for cross-kingdom transfers between fungi and non-fungal Holomycota, as well as between deeply divergent protists and NDF. The propensity for transfers between kingdoms could be explained by shared lifestyles, as some non-Dikarya clades share key traits with protists, such as thriving in aquatic environments and retaining flagella for motility^3,13^. Taken together, our study provides the first large-scale insights into the evolutionary origins of the EDF mobilome, its impacts on genome regulation and the emergence of new functions. Insights from EDF contribute significantly to the emerging view that TEs are not merely genomic parasites, but rather foundational drivers of structural plasticity and evolutionary innovation. Elucidating such key mechanisms of genome evolution helps to recapitulate the highly dynamic nature of fungal genome architecture.

## METHODS

### Species tree reconstruction

Proteomes from 238 fungal and non-fungal species were retrieved from JGI or NCBI (Supplementary File 1). The dataset includes 163 genomes from NDF with at least one species for each genus with available data, 27 Dikarya genomes (Ascomycota and Basidiomycota, from different clades representing the diversity of these groups), along with 48 genomes covering non-fungal species (non-fungal Holomycota and Amoebozoa representatives). From annotated proteomes for each genome, single-copy orthogroups (1) were identified with OrthoFinder (v3.1)^63^. The marker set was filtered using PhyKit (v2.0)^64^ to retain only groups with taxon occupancy ≥ 40 % (2), post-trimming alignment length ≥ 60 amino acid columns (3), parsimony-informative sites (PIS, counted from trimmed alignment) ≥ 25 (4) and relative composition variability (RCV) ≤ 0.5 (5). Pergene trees (IQ-TREE) (v2.3.6)^65^ (--fast mode) were used to detect long-branch outlier sequences: any taxon whose terminal branch length exceeded the per-gene mean by more than 3.5 standard deviations was flagged as a potential paralog, removed, and the gene was re-evaluated against criteria 1–5. A total of 682 markers passed all filters, yielding a concatenated supermatrix of 238 taxa × 219,476 amino acid positions. Each retained marker was individually aligned with MAFFT (v7.520)^66^ (--auto; for alignments ≥ 200 sequences, --retree 2 --reorder was applied to reduce computational load). Alignments were trimmed with trimAl (v1.4.1)^67^, removing columns with more than 20% gaps. The trimmed alignments were concatenated and partitioned by gene. A first-pass guide tree (- B 1000) was computed to seed the final analysis. The final tree was inferred under the LG+G4 model. Licenses for silhouettes included in Figure 1 are listed in Supplementary File 8.

### TE annotation

TEs were annotated in all genomes using EarlGrey (v7)^68^. The fungal-specific curated TE library, Mycomobilome^36^, was combined with de novo annotation of non-fungal genomes following a similar approach. Briefly, consensus sequences were generated using - earlGreyLibConstruct with Dfam 3.7 curated elements as the reference database, using default settings. All per-species consensus sequences were combined into a single multi-FASTA file and clustered with a scalable cascaded clustering approach using MMseqs2 (v14.7564)^69^ with --min-seq-id 0.8 -c 0.8 --cov-mode 1 --cluster-reassign, retaining the representative sequence for each cluster. Sequences shorter than 80 bp and with > 50 % simple or low-complexity sequence were discarded. To identify and remove putative host genes that may have been captured in the TE library, open reading frames were identified in all consensus sequences and matched against the RefSeq non-redundant protein database using DIAMOND BLASTX v2.1.8^70^. Sequences with a RefSeq protein hit (identity ≥ 40 %, e-value ≤ 10⁻⁵) that did not also match known TE proteins (i.e. the best RefSeq hit annotation did not contain the terms transposon, transposase, integrase, reverse transcriptase, capsid, or gag) were classified as host-derived and removed from the TE library. Each consensus was classified against the Dfam 3.7 database (20,462 HMM profiles) using nhmmer (HMMER 3.3.2). TE superfamily nomenclature follows Wicker et al. (2007)^71^. The new combined library was used for annotation of all species. Two Microsporidia genomes (MI_Encro1, MI_Ordco1) returned zero TE content after filtering and were excluded from ordination analyses but retained in all other comparative analyses. Kimura divergence scores were obtained from EarlGrey outputs and used to calculate the recency index (RI) as the proportion of the total TE content (measured in base pairs) that consists of recently inserted copies. Lengths of all annotated TE copies with K2P distance < 0.10 were summed and divided by the total length of all annotated TE copies in the genome. PCA of TE composition was conducted using eight high-level classes (DNA, LTR, LINE, SINE, Helitron, RC, PLE, Unknown) as proportions summing to 1. The Shannon diversity index (H’) measures the overall complexity of the TE community. It is calculated as: for each superfamily, multiply its proportional contribution to total TE base pairs by the log₂ of that proportion, sum these products across all superfamilies, and negate the result. A value of H′ = 0 bits means the entire TE content is dominated by a single superfamily; higher values indicate a more diverse community.

### Comparative genomics, pangenome and synteny analyses

Genome size, gene density, and intron density were calculated from genomic FASTA files and GFF annotation files for each species. InterProScan (v5.67-99.0)^72^ was used to generate domain annotation for each species, including domains of unknown function. Pan-genome accumulation curves were generated as in Steindorff et al. (2024)^73^. Briefly, 100 random permutations of species-addition order were applied to the OrthoFinder gene presence/absence matrix. At each depth (number of species sampled), orthogroups were classified as core (present in all sampled species), shell (present in ≥ 2 but < n genomes), or unique/cloud (present in exactly one sampled species). The mean and standard deviation across 100 random orderings were calculated per depth for each clade. Microsynteny conservation across 17 clades of the NDF dataset was estimated from the frequency of collinear ortholog pairs, following the approach of Grau-Bové et al. (2017)^74^. Gene positions were extracted from GFF/GFF3 annotation files for each species. Per scaffold, protein-coding gene features were sorted by chromosomal position using the locus_tag attribute (NCBI-sourced genomes) or the Name/ID attribute (JGI-sourced genomes) as the gene identifier. Scaffolds with fewer than three annotated genes were excluded. For each species pair (i, j) within the same clade, collinear ortholog pairs were identified as follows. On each scaffold of species i, the ordered sequence of orthologs shared with species j was extracted. For each consecutive pair of orthologs on the same scaffold of species i, we tested whether their corresponding orthologs in species j were also adjacent on the same scaffold, irrespective of gene orientation. To account for spurious gene-order conservation from random co-localization, random genomic positions were assigned to each gene using bedtools shuffle (v2.24)^75^ in 100 independent rounds per species, with scaffolds preserved as location constraints (-chrom flag). Each replicate used a fixed per-replicate random seed. The mean over 100 replicates was computed for each species pair. Synteny score ratios were corrected against patristic distances extracted from the species tree, computed as the sum of branch lengths along the path connecting the two tips. Syntenic gene-pair links were visualized using Circos (v0.67)^76^. For nucleosome occupancy prediction, we used the fungal-trained algorithm Nuclpred^42^. Occupancy and the autocorrelation value identifying nucleosome phasing and regularity were computed from the generated output WIG files. For in vivo nucleosome occupancy data, we used MNase-seq data from *R. microsporus*, *Neurospora crassa* and *Botrytis cinerea* (Supplementary File 10). Reads were aligned to reference genomes using Bowtie2 (v2.5.1)^77^. BAMs were coordinate-sorted with samtools (v1.17)^78^ and PCR duplicates were removed with Picard MarkDuplicates (v3.1). DANPOS (v3)^79^ (dpos subcommand) was applied to the pooled BAM to call per-nucleosome dyad positions and compute per-nucleosome metrics. The output positions.xls file provides, for every called nucleosome, the smoothed dyad position (smt_pos), peak height (smt_value, proportional to nucleosome sharpness/occupancy), and fuzziness score (standard deviation of the read-end distribution around the dyad, in bp; lower values indicate better-positioned nucleosomes). A smoothed WIG occupancy file (*.smooth.wig) was also produced for regional analyses. TEs in euchromatin (H3K9me3-negative) were identified in *R. microsporus.* For each TE ≥500 bp with ≥2 nucleosomes, a dyad density profile was built with 1 kb flanks (50 linear bins) and a TE body scaled to 100 bins. Occupancy profile fuzziness (1 − Pearson correlation between each TE’s profile and its group mean) was computed to quantify inter-TE nucleosome heterogeneity. Differences between MAC and non-MAC groups were assessed using the Kolmogorov-Smirnov test.

### TE-associated domain expansion

To identify protein domains that have undergone clade-specific expansions in parallel with TE activity, a systematic scan of all Pfam domains was performed across the species IPR.tab dataset. For each Pfam domain with ≥ 5 total protein hits across the dataset, two expansion tests were applied: (i) Clade expansion: a one-sided Mann–Whitney U test comparing per-species copy counts in each clade against all other species. A domain was flagged as clade expanded if the corrected p-value was < 10⁻⁴ (BH-FDR < 0.05 across all clades × domain combinations), the fold-change over the rest-of-dataset median was ≥ 5, and the within-clade median copy count was ≥ 3 copies per species. (ii) Species-specific expansion: a domain was flagged if any single species harbored ≥ 10 copies, while the median of all other species was ≤ 1 copy. Ubiquitous housekeeping domains (present in > 40 % of species with ≥ 2 copies), ribosomal and proteasomal components were excluded. For each species with at least one gene carrying an expanded domain, the distance from every such gene to the nearest annotated TE copy was computed with bedtools closest^75^. Three comparisons were made per domain × species: (A) expanded-clade species vs. non-expanded-clade species for the same domain; (B) target-domain genes vs. a size-matched random background of other genes in the same genome; (C) high-copy species (top 25 % by domain copy number) vs. low-copy species (bottom 75 %) within the expanded clade. Background genes were sampled without replacement from the same genome. Two-sided Mann–Whitney U tests with BH correction were applied. For domains with significant TE proximity and TE-content correlation, structural analysis was performed to identify putative pack-TEs. Each candidate gene was intersected with the TE annotation; genes fully embedded within a TE interval (overlap ≥ 90 % of gene length by a single TE copy) and shorter than the median domain-family gene length by > 40 % were flagged as putative pack elements.

### Epigenetics data analysis

DNMT and DIM2 DNA methyltransferases were identified in proteomes using InterProScan v5.67-99.0 (Pfam 36.0 database)^72^. For each protein, the presence of the following Pfam domains was recorded: C5MT catalytic domain (PF00145), RFTS replication foci targeting sequence, BAH bromo-adjacent homology, DNAmet_Nter N-terminal methyltransferase extension, Helicase_C, SNF2_N, and PHD finger. Reciprocal BLASTP (BLAST+ 2.12.0; E ≤ 10⁻⁵) was performed against a curated set of reference DNMT1, DIM-2, and RID/DNMT2 sequences from characterized fungal species. Final candidates were included in Supplementary File 4. For 5mC methylation analyses, the methylation score was computed as the weighted methylation ratio (methylated reads/methylated + unmethylated reads)^18^ over each TE. For H3K9me3 data, reads were independently quality-trimmed with Trim Galore (v0.6.10) and aligned to the reference genome with Bowtie2 (v2.5.1)^77^. Peaks were called with MACS2 (v2.2.9.1)^80^, using the input-control BAM for background correction. H3K9me3 enrichment per TE copy was computed as the mean bedgraph signal over the copy coordinates, normalized to RPKM. For sRNA data processing, small RNA (sRNA) sequencing reads were adapter-trimmed with Cutadapt (v4.6) --minimum-length 18 --maximum-length 32. Trimmed reads were size-selected to 20–25 nt. Size-selected reads were mapped to the cross-species TE consensus library with Bowtie2 (v2.5.1)^77^. Reads per million (RPM) were computed per TE family. 6mA data were processed as previously described. Raw reads were filtered using SFilter to remove short reads and reads derived from sequencing adapters. Filtered reads were aligned to the reference genome using BLASR (v1.5.0)^81^. Modified sites were then identified through kinetic analysis of the aligned DNA sequence data. Detected 6mA sites were further filtered with a minimum 15x per-strand coverage to remove potential false positives and select only 6mA sites with >25 mQv. 6mA clusters were defined as previously established^19,60^. MACs overlapping the TE transcription start site window (−150 to +400 bp) were used for the MAC presence-absence analysis. Gene-level 6mA density was computed as the number of methylated sites per kilobase of feature body. Average 6mA profiles over genomic regions were computed with deepTools (v2)^82^.

### TE domain co-option and TF binding site enrichment in TE sequences

Annotated InterProScan proteomes were used to classify genes on the basis of their Pfam domain composition, using a curated list of TE-machinery Pfam accessions (Supplementary File 9)^44,83^. A gene was designated co-opted if it satisfied both of the following: (i) at least one te core or te_assoc domain is present; and (ii) the gene is not classified as a complete TE ORF, i.e. it carries at least one Pfam domain not belonging to the TE domain set. A total of 51,799 co-opted genes were identified across 238 species. Chimeric transcripts were detected using ChimeraTE (Mode 1, RNA-based)^84^. For precise characterization of TE fusion of expanded domains in NDF, Pfam domain accessions were collected from the IPR.tab file. To assess the association between expanded domains and TE fusions, TE libraries for all 238 species were first translated into all six reading frames with EMBOSS getorf (minimum ORF length 100 aa). The resulting ORF databases were then screened by hmmscan against a curated set of HMMs: FLYWCH (PF04500), CotH (PF08757) and NACHT (PF05729), retaining hits at E-value ≤ 10⁻⁵. For the co-opted domains, protein sequences from the 238 proteomes were extracted and aligned with MAFFT (v7.526)^66^. Alignments were trimmed with trimAl (v1.5)^67^ and a maximum-likelihood tree was inferred with FastTree (v2.2)^85^ (-lg -gamma). Each leaf was annotated with the clade identity of its host species and flagged if the protein also carried any TE-machinery domain. To assess the full breadth of methyltransferase domains encoded within TE consensus sequences, a similar process was conducted for C5-cytosine DNA methyltransferase (DNMT): PF00145 (DNA_methylase), PF17980 (ADD_DNMT3), (MTA70): PF05063 (MT-A70) and DAM-class N6-adenine DNA methyltransferase: PF05869, and PF01555 (N6_N4_Mtase). A TE ORF was classified as carrying a given methyltransferase domain when ≥ 1 hmmscan hit at E-value ≤ 10⁻⁵ (domain-level score) was detected. Only DAM-domain ORFs also carried TE-structural domains in the same ORF (co-occurrence with RVT_1, rve, Chromo, or gag_pre-integrase at E ≤ 10⁻⁵). Reverse transcriptase (RT) and DAM domain trees were generated as described above.

For TF binding analysis, DAP-seq peak data for 58 transcription factors from *R. microsporus* ATCC11559 were considered. DAP-seq peak coordinates were intersected with TE annotation using bedtools intersect. For each TF, the fraction of peaks overlapping any TE copy was computed; enrichment relative to the genomic TE coverage was tested with a two-sided Fisher’s exact test. Significance was assessed after Benjamini–Hochberg correction (FDR 5 %). FIMO motif enrichment in TE sequences (script 03). FIMO (v5.5.5)^86^ was run for each of the 58 TF motif files against the TE sequences of each TE class, and separately against 3× size-matched genomic shuffled sequences generated with bedtools shuffle. The p-value threshold for FIMO hit reporting was 1 × 10⁻⁴. Enrichment in TE sequences relative to the shuffled background was tested with a two-sided Fisher’s exact test on the 2 × 2 contingency table (TE sequences with ≥ 1 FIMO hit vs. background sequences with ≥ 1 FIMO hit); p-values were corrected across all TF × TE class combinations using the Benjamini–Hochberg procedure (FDR 5 %). For motif prediction in GremLINE elements, the 44 identified sequences in *M. lusitanicus* were scanned with FIMO following the same procedure and the same e-value filtering thresholds. For the classification of Gypsy elements recognized by HLH TFs, the RT domain sequences of all 103 Gypsy elements in *R. microsporus* ATCC11559 carrying an identifiable RT domain were extracted and aligned. A maximum-likelihood tree was inferred with IQ-TREE2 (-m MFP -bb 1000 --redo). Gypsy elements were classified into RT-domain clades using TEsorter (v1.4.7)^87^ with the REXdb/Metazoa database (--db REXdb -p 4), which resolved four clades: Chlamyvir, Reina, chromo-outgroup, and non-chromo-outgroup.

### Whole genome duplication analysis

For WGD analyses, only MU genomes with fewer than 200 scaffolds were considered. Whole-proteome all-vs-all protein similarity was computed for each selected species with wgd dmd (wgd v2)^88^, followed by clustering of paralogous gene families using the MCL algorithm (--inflation 1.5). For each paralogous pair, the coding sequences were aligned with PAL2NAL (v14)^89^. Ks and Ka values were estimated from the resulting codon alignment with yn00 PAML (v4.10.10). Gene pairs were excluded if: Ks = 0 (CDS identity; likely assembly duplicates), Ks > 3.0 (saturation under any standard substitution rate), Ka/Ks > 5 (non-synonymous model convergence failure), or the codon alignment contained fewer than 100 codons after gap removal. Ks distributions were modeled using Gaussian mixture models, with the optimal number of components selected by Bayesian Information Criterion (BIC), and putative WGD peaks were defined as components with weight ≥ 0.10 and 0.05 < Ks < 3.0. Intra-genomic collinearity was assessed using wgd syn (wgd v2), which builds a collinearity graph using the MCScan-derived algorithm: it first identifies anchor gene pairs and then clusters anchors into collinear blocks by the tandem-chaining heuristic. Each collinear block is reported as a multiplicon: a pair of genomic segments on the same or different chromosomes. Branch calibration was computed based on a fossil-calibrated tree of fungi^16^. The boundaries of intra-genomic collinear blocks were used to define breakpoint windows (10 kb). A breakpoint was defined as the chromosomal interval between the last anchor-point coordinate of one multiplicon and the first anchor-point coordinate of the next multiplicon on the same scaffold, sorted by genomic position. One hundred randomly sampled windows of the same length distribution were used as background. A two-sided Wilcoxon signed-rank test compared matched breakpoint vs. random TE densities, one observation per breakpoint, paired. TE insertion ages were estimated using Kimura divergence and converted into absolute time (Mya) using lineage-specific substitution rates following t = K/(2r). WGD ages were estimated from Ks peak positions using the same calibration framework. TE bursts were classified relative to WGD timing as pre-WGD, co-WGD (within a twofold temporal window), or post-WGD, and associations between TE and WGD timing were evaluated using Spearman correlation.

TE copy number evolution was assessed using species × superfamily matrices, comparing WGD and non-WGD species using Mann–Whitney U tests. Log2 fold enrichment was computed as log2((mean_WGD + 1)/(mean_nonWGD + 1)). Finally, phylogenetic relationships, WGD calls, TE genomic content, and synteny multiplicon counts were integrated into a comparative framework to summarize genome-wide evolutionary patterns across species.

### Assessment of horizontal transposon transfers

HTT detection was performed based on previous approaches developed for insect and fungal TE landscapes^55,90^. To calibrate evolutionary divergence between each pair of clades at the node level, the synonymous substitution rate (Ks) of core single-copy genes was computed for each internal node of the species tree. For each node, single-copy orthogroups common to both descendant sub-clades were used as the gene set. The two CDS were aligned at the protein level with MAFFT (v7.520)^66^, then a codon alignment was built with PAL2NAL (v14)^89^. Ks values were estimated with the kaks function in seqinr (v4.2-36)^91^. The 0.5th percentile of the Ks distribution per node was used as a node Ks threshold. TE copies < 300 bp were discarded and all retained TE sequences were extracted from genome FASTAs with BEDtools (v2.31.1) getfasta and compiled into a combined FASTA. To restrict HTT analysis to copies likely to encode functional TE proteins, TE-related protein domains were identified in each copy. TE sequences were translated in all six frames with EMBOSS transeq (v6.6.0). Translated sequences were searched against: (i) 142 Conserved Domain Database (CDD) profiles with RPSTBLASTN (BLAST+ v2.12.0, -evalue 0.001 -max_target_seqs 1); (ii) 110 TE-related Pfam HMM profiles with HMMER v3.4 (hmmsearch -E 0.001). Only TE copies with at least one domain hit passing the e-value threshold were retained for the HTT analysis. Retained domain-containing TE sequences were searched against each other in an all-vs-all BLASTN (BLAST+ v2.12.0)^92^. Cross-clade hits (i.e. query and subject from different clades) were retained if percent identity ≥ 75 %, alignment length ≥ 300 bp, and bitscore ≥ 200. Fragmented collinear hits between the same pair of TE sequences were chained into single extended alignments, allowing a maximum gap and maximum overlap of 600 bp. Pairs were further filtered to require that the chained alignment covered ≥ 60 % of both the query and subject TE length (to exclude partial matches that could reflect conserved domain fragments). This yielded 50,143 chained cross-clade hit pairs as input to Ks estimation. A candidate HTT pair was retained if its estimated Ks was lower than the 0.5th-percentile Ks threshold of the most recent common ancestor (MRCA) node of the two species’ clades (i.e. the TE copies are more similar than expected for any gene evolving neutrally since the two lineages diverged). This yielded 223 HTT candidate pairs spanning 20 clade pairs. Because multiple HTT pairs stemming from the same biological event inflate the candidate count, HTT pairs were clustered into independent transfer communities. For each pair of clades sharing at least one HTT candidate, an intra-clade all-vs-all BLASTN was performed (-dbsize 10,000,000,000 -evalue 10 - max_target_seqs 600,000) to obtain within-clade identity scores. HTT pairs were represented as nodes in a graph; two nodes were connected by an edge if: (a) they involved the same TE copy, or (b) the between-clade identity of the TE pair exceeded the within-clade identity of the same TE (i.e. the cross-species copy is more similar than copies within the same species, ruling out independent parallel evolution). The Louvain community detection algorithm (igraph v0.11.8) was applied to identify communities. Communities with inter-community edge density > 0.05 were merged to minimize false positives. This yielded 20 HTT communities across the studied clades. To account for the possibility that a single ancestral transfer produced multiple descendant lineages that each appear as independent transfers, a second-level network analysis was applied. Communities were treated as nodes; communities sharing TE copies were connected. Single-linkage clustering with NetworkX (v2.5.1) identified clusters of related communities. For each cluster, the species tree was pruned to include only the species and collapsed nodes present in that cluster using ete3 (v3.1.3)^93^. Per node of the pruned tree, the minimum number of independent transfers required to explain the observed pattern parsimoniously was counted: if multiple transfers from different clade pairs involved the same TE community, one event sufficed; if the same clade pair was involved in multiple independent communities, that exact count was recorded. Summing across all nodes yielded a minimum of 15 independent HTT events.

## ACKNOWLEDGMENTS

We thank Felix Zimmermann for insightful discussions about horizontal transposon transfers and the rest of the group members for feedback and discussions.

## CODE AVAILABILITY

Code used for this work is available at: https://doi.org/10.5281/zenodo.24369142

## AUTHOR CONTRIBUTIONS

C.L., G.T. and D.C. conceived the study. C.L. and G.T. analyzed the data. C.L. wrote the manuscript with significant input from G.T. and D.C. D.C. acquired funding and supervised the study.

## REFERENCES

1. Case, N. T. et al. Fungal impacts on Earth’s ecosystems. Nature 638, 49–57 (2025).

2. Heitman, J. The Fungal Kingdom as a Rosetta Stone for biological discovery. Curr. Biol. 35, R427–R433 (2025).

3. Amses, K. R. et al. Diploid-dominant life cycles characterize the early evolution of Fungi. Proc. Natl. Acad. Sci. U. S. A. 119, e2116841119 (2022).

4. James, T. Y., Stajich, J. E., Hittinger, C. T. & Rokas, A. Toward a Fully Resolved Fungal Tree of Life. Annu. Rev. Microbiol. 74, 291–313 (2020).

5. James, T. Y. & Rokas, A. Use their names: there are no basal, lower, or early diverging fungi. Mycologia 117, 246–254 (2025).

6. Prostak, S. M., Robinson, K. A., Titus, M. A. & Fritz-Laylin, L. K. The actin networks of chytrid fungi reveal evolutionary loss of cytoskeletal complexity in the fungal kingdom. Curr. Biol. 31, 1192–1205.e6 (2021).

7. Weete, J. D., Abril, M. & Blackwell, M. Phylogenetic Distribution of Fungal Sterols. PLoS ONE 5, e10899 (2010).

8. Orłowska, M., Steczkiewicz, K. & Muszewska, A. Utilization of Cobalamin Is Ubiquitous in Early-Branching Fungal Phyla. Genome Biol. Evol. 13, evab043 (2021).

9. Medina, E. M., Turner, J. J., Gordân, R., Skotheim, J. M. & Buchler, N. E. Punctuated evolution and transitional hybrid network in an ancestral cell cycle of fungi. eLife 5, e09492 (2016).

10. Galindo, L. J., López-García, P., Torruella, G., Karpov, S. & Moreira, D. Phylogenomics of a new fungal phylum reveals multiple waves of reductive evolution across Holomycota. Nat. Commun. 12, 4973 (2021).

11. Torruella, G. et al. Global transcriptome analysis of the aphelid *Paraphelidium tribonemae* supports the phagotrophic origin of fungi. *Commun*. Biol. 1, 231 (2018).

12. Chang, Y. et al. Genome-scale phylogenetic analyses confirm *Olpidium* as the closest living zoosporic fungus to the non-flagellated, terrestrial fungi. Sci. Rep. 11, 3217 (2021).

13. Berbee, M. L., James, T. Y. & Strullu-Derrien, C. Early Diverging Fungi: Diversity and Impact at the Dawn of Terrestrial Life. Annu. Rev. Microbiol. 71, 41–60 (2017).

14. Voigt, K. et al. Early-diverging fungal phyla: taxonomy, species concept, ecology, distribution, anthropogenic impact, and novel phylogenetic proposals. Fungal Divers. 109, 59–98 (2021).

15. Liu, H. et al. A taxon-rich and genome-scale phylogeny of Opisthokonta. PLOS Biol. 22, e3002794 (2024).

16. Szánthó, L. L. et al. A timetree of Fungi dated with fossils and horizontal gene transfers. *Nat*. Ecol. Evol. 1–13 (2025) doi:10.1038/s41559-025-02851-z.

17. Merényi, Z. et al. Genomes of fungi and relatives reveal delayed loss of ancestral gene families and evolution of key fungal traits. *Nat*. Ecol. Evol. 7, 1221–1231 (2023).

18. Bewick, A. J. et al. Diversity of cytosine methylation across the fungal tree of life. *Nat*. Ecol. Evol. 3, 479–490 (2019).

19. Lax, C. et al. Symmetric and asymmetric DNA N6-adenine methylation regulates different biological responses in Mucorales. Nat. Commun. 15, 6066 (2024).

20. Mondo, S. J. & Grigoriev, I. V. A genomic perspective on fungal diversity and evolution. Nat. Rev. Microbiol. 23, 718–733 (2025).

21. Badet, T. & Croll, D. Phylogenomic signatures of repeat-induced point mutations across the fungal kingdom. PLOS Biol. 23, e3003433 (2025).

22. Muszewska, A., Steczkiewicz, K., Stepniewska-Dziubinska, M. & Ginalski, K. Transposable elements contribute to fungal genes and impact fungal lifestyle. Sci. Rep. 9, 4307 (2019).

23. Gusa, A. et al. Transposon mobilization in the human fungal pathogen *Cryptococcus* is mutagenic during infection and promotes drug resistance in vitro. Proc. Natl. Acad. Sci. 117, 9973–9980 (2020).

24. Mackey, A. I. et al. Temperature and genetic background drive mobilization of diverse transposable elements in a global human fungal pathogen. PLOS Genet. 22, e1011979 (2026).

25. Badet, T., Tralamazza, S. M., Feurtey, A. & Croll, D. Recent reactivation of a pathogenicity-associated transposable element is associated with major chromosomal rearrangements in a fungal wheat pathogen. Nucleic Acids Res. 52, 1226–1242 (2023).

26. Tahiri, G., Lax, C., Nicolás, F. E., Garre, V. & Navarro, E. Direct Targeted Degradation of Transposon RNAs by the Non-Canonical RNAi Pathway of the Fungus *Mucor lusitanicu*s. Int. J. Mol. Sci. 26, 2738 (2025).

27. Haase, M. A. B. et al. Ancient co-option of LTR retrotransposons as yeast centromeres. Nature 651, 1004–1011 (2026).

28. Gluck-Thaler, E. et al. Giant *Starship* Elements Mobilize Accessory Genes in Fungal Genomes. Mol. Biol. Evol. 39, msac109 (2022).

29. Sato, Y. et al. Starship giant transposons dominate plastic genomic regions in a fungal plant pathogen and drive virulence evolution. Nat. Commun. 16, 6806 (2025).

30. Urquhart, A. S., Vogan, A. A., Gardiner, D. M. & Idnurm, A. *Starships* are active eukaryotic transposable elements mobilized by a new family of tyrosine recombinases. Proc. Natl. Acad. Sci. 120, e2214521120 (2023).

31. Grigoriev, I. V. et al. MycoCosm portal: gearing up for 1000 fungal genomes. Nucleic Acids Res. 42, D699–D704 (2014).

32. Marcet-Houben, M. & Gabaldón, T. Beyond the Whole-Genome Duplication: Phylogenetic Evidence for an Ancient Interspecies Hybridization in the Baker’s Yeast Lineage. PLOS Biol. 13, e1002220 (2015).

33. Corrochano, L. M. et al. Expansion of Signal Transduction Pathways in Fungi by Extensive Genome Duplication. Curr. Biol. 26, 1577–1584 (2016).

34. Martin, F. et al. The genome of *Laccaria bicolor* provides insights into mycorrhizal symbiosis. Nature 452, 88–92 (2008).

35. Yang, P., Wang, D. & Kang, L. Alternative splicing level related to intron size and organism complexity. BMC Genomics 22, 853 (2021).

36. Baril, T. & Croll, D. MycoMobilome: a community-focused non-redundant database of transposable element consensus sequences for the fungal kingdom. NAR Genomics Bioinforma. 8, lqag026 (2026).

37. Baril, T., Puccetti, G. & Croll, D. Historic transposon mobilisation waves create distinct pools of adaptive variants in a major crop pathogen. Nat. Commun. 16, 9961 (2025).

38. Idnurm, A., Walton, F. J., Floyd, A. & Heitman, J. Identification of the sex genes in an early diverged fungus. Nature 451, 193–196 (2008).

39. Nikitin, D. M. Transposable element-host genome evolutionary arms race revealed by multi-modal epigenomic profiling in a telomere-to-telomere human genome reference. R. Soc. Open Sci. 13, rsos260639 (2026).

40. Lax, C. et al. Symmetric adenine methylation is an essential DNA modification in the early-diverging fungus *Rhizopus microsporus*. Nat. Commun. 16, 3843 (2025).

41. Romero Charria, P., et al. Adenine DNA methylation associated with transcriptionally permissive chromatin is widespread across eukaryotes. Nat. Genet. 57, 3126–3136 (2025).

42. Mondo, S. J. et al. Consecutive low-frequency shifts in A/T content denote nucleosome positions across microeukaryotes. iScience 28, 112472 (2025).

43. Trieu, T. A. et al. A Non-canonical RNA Silencing Pathway Promotes mRNA Degradation in Basal Fungi. PLOS Genet. 11, e1005168 (2015).

44. Oggenfuss, U., Badet, T. & Croll, D. A systematic screen for co-option of transposable elements across the fungal kingdom. Mob. DNA 15, 2 (2024).

45. Szebenyi, C. et al. cotH Genes Are Necessary for Normal Spore Formation and Virulence in *Mucor lusitanicus*. mBio 14, e03386–22 (2023).

46. Gebremariam, T. et al. Anti-CotH3 antibodies protect mice from mucormycosis by prevention of invasion and augmenting opsonophagocytosis. Sci. Adv. 5, eaaw1327 (2019).

47. Chibucos, M. C. et al. An integrated genomic and transcriptomic survey of mucormycosis-causing fungi. Nat. Commun. 7, 12218 (2016).

48. Fernández-Moreno, M. et al. Pervasive co-option of prokaryotic adenine methyltransferases by eukaryotic retrotransposons. Preprint at 10.64898/2026.05.21.726589 (2026).

49. Lax, C. et al. Unveiling a pervasive DNA adenine methylation regulatory network in the early-diverging fungus *Rhizopus microsporus*. Nat. Commun. 16, 10277 (2025).

50. Navarro-Mendoza, M. I. et al. Early Diverging Fungus *Mucor circinelloides* Lacks Centromeric Histone CENP-A and Displays a Mosaic of Point and Regional Centromeres. Curr. Biol. 29, 3791–3802.e6 (2019).

51. Ma, L.-J. et al. Genomic Analysis of the Basal Lineage Fungus *Rhizopus oryzae* Reveals a Whole-Genome Duplication. PLoS Genet. 5, e1000549 (2009).

52. Gryganskyi, A. P. et al. Phylogenetic and Phylogenomic Definition of *Rhizopus* Species. G3 Genes Genomes Genet. 8, g3.200235.2018 (2018).

53. David, K. T. et al. Discovery of additional ancient genome duplications in yeasts. Curr. Biol. 36, 1067–1075.e3 (2026).

54. Corrochano, L. M. et al. Expansion of Signal Transduction Pathways in Fungi by Extensive Genome Duplication. Curr. Biol. 26, 1577–1584 (2016).

55. Romeijn, J., Bañales, I. & Seidl, M. F. Extensive horizontal transfer of transposable elements shapes fungal mobilomes. Curr. Biol. 10.1016/j.cub.2025.11.012 (2025) doi:10.1016/j.cub.2025.11.012.

56. Wang, Y. et al. Divergent Evolution of Early Terrestrial Fungi Reveals the Evolution of Mucormycosis Pathogenicity Factors. Genome Biol. Evol. 15, evad046 (2023).

57. De Souza, F. S. J., Franchini, L. F. & Rubinstein, M. Exaptation of Transposable Elements into Novel Cis-Regulatory Elements: Is the Evidence Always Strong? Mol. Biol. Evol. 30, 1239–1251 (2013).

58. Slotkin, R. K. & Martienssen, R. Transposable elements and the epigenetic regulation of the genome. Nat. Rev. Genet. 8, 272–285 (2007).

59. Lev, I., Gingold, H. & Rechavi, O. H3K9me3 is required for inheritance of small RNAs that target a unique subset of newly evolved genes. eLife 8, e40448 (2019).

60. Mondo, S. J. et al. Widespread adenine N6-methylation of active genes in fungi. Nat. Genet. 49, 964–968 (2017).

61. Kellis, M., Birren, B. W. & Lander, E. S. Proof and evolutionary analysis of ancient genome duplication in the yeast Saccharomyces cerevisiae. Nature 428, 617–624 (2004).

62. Wolfe, K. H. & Shields, D. C. Molecular evidence for an ancient duplication of the entire yeast genome. Nature 387, 708–713 (1997).

63. Emms, D. M. & Kelly, S. OrthoFinder: phylogenetic orthology inference for comparative genomics. Genome Biol. 20, 238 (2019).

64. Steenwyk, J. L. et al. PhyKIT: a broadly applicable UNIX shell toolkit for processing and analyzing phylogenomic data. Bioinformatics 37, 2325–2331 (2021).

65. Nguyen, L.-T., Schmidt, H. A., Von Haeseler, A. & Minh, B. Q. IQ-TREE: A Fast and Effective Stochastic Algorithm for Estimating Maximum-Likelihood Phylogenies. Mol. Biol. Evol. 32, 268–274 (2015).

66. Katoh, K. & Standley, D. M. MAFFT Multiple Sequence Alignment Software Version 7: Improvements in Performance and Usability. Mol. Biol. Evol. 30, 772–780 (2013).

67. Capella-Gutiérrez, S., Silla-Martínez, J. M. & Gabaldón, T. trimAl: a tool for automated alignment trimming in large-scale phylogenetic analyses. Bioinformatics 25, 1972–1973 (2009).

68. Baril, T., Galbraith, J. & Hayward, A. Earl Grey: A Fully Automated User-Friendly Transposable Element Annotation and Analysis Pipeline. Mol. Biol. Evol. 41, msae068 (2024).

69. Steinegger, M. & Söding, J. MMseqs2 enables sensitive protein sequence searching for the analysis of massive data sets. Nat. Biotechnol. 35, 1026–1028 (2017).

70. Buchfink, B., Xie, C. & Huson, D. H. Fast and sensitive protein alignment using DIAMOND. Nat. Methods 12, 59–60 (2015).

71. Wicker, T. et al. A unified classification system for eukaryotic transposable elements. Nat. Rev. Genet. 8, 973–982 (2007).

72. Jones, P. et al. InterProScan 5: genome-scale protein function classification. Bioinformatics 30, 1236–1240 (2014).

73. Steindorff, A. S. et al. Comparative genomic analysis of thermophilic fungi reveals convergent evolutionary adaptations and gene losses. *Commun*. Biol. 7, 1124 (2024).

74. Grau-Bové, X. et al. Dynamics of genomic innovation in the unicellular ancestry of animals. eLife 6, e26036 (2017).

75. Quinlan, A. R. & Hall, I. M. BEDTools: a flexible suite of utilities for comparing genomic features. Bioinformatics 26, 841–842 (2010).

76. Krzywinski, M. et al. Circos: An information aesthetic for comparative genomics. Genome Res. 19, 1639–1645 (2009).

77. Langmead, B. & Salzberg, S. L. Fast gapped-read alignment with Bowtie 2. Nat. Methods 9, 357–359 (2012).

78. Li, H. et al. The Sequence Alignment/Map format and SAMtools. Bioinformatics 25, 2078–2079 (2009).

79. Chen, K. et al. DANPOS: Dynamic analysis of nucleosome position and occupancy by sequencing. Genome Res. 23, 341–351 (2013).

80. Feng, J., Liu, T., Qin, B., Zhang, Y. & Liu, X. S. Identifying ChIP-seq enrichment using MACS. Nat. Protoc. 7, 1728–1740 (2012).

81. Chaisson, M. J. & Tesler, G. Mapping single molecule sequencing reads using basic local alignment with successive refinement (BLASR): application and theory. BMC Bioinformatics 13, 238 (2012).

82. Ramírez, F. et al. deepTools2: a next generation web server for deep-sequencing data analysis. Nucleic Acids Res. 44, W160–W165 (2016).

83. Grau-Bové, X. et al. A phylogenetic and proteomic reconstruction of eukaryotic chromatin evolution. *Nat*. Ecol. Evol. 6, 1007–1023 (2022).

84. Oliveira, D. S. et al. ChimeraTE: a pipeline to detect chimeric transcripts derived from genes and transposable elements. Nucleic Acids Res. 51, 9764–9784 (2023).

85. Price, M. N., Dehal, P. S. & Arkin, A. P. FastTree 2 – Approximately Maximum-Likelihood Trees for Large Alignments. PLoS ONE 5, e9490 (2010).

86. Bailey, T. L., Johnson, J., Grant, C. E. & Noble, W. S. The MEME Suite. Nucleic Acids Res. 43, W39–W49 (2015).

87. Zhang, R.-G. et al. TEsorter: An accurate and fast method to classify LTR-retrotransposons in plant genomes. Hortic. Res. 9, uhac017 (2022).

88. Chen, H., Zwaenepoel, A. & Van De Peer, Y. wgd v2: a suite of tools to uncover and date ancient polyploidy and whole-genome duplication. Bioinformatics 40, btae272 (2024).

89. Suyama, M., Torrents, D. & Bork, P. PAL2NAL: robust conversion of protein sequence alignments into the corresponding codon alignments. Nucleic Acids Res. 34, W609–W612 (2006).

90. Zhang, H.-H., Peccoud, J., Xu, M.-R.-X., Zhang, X.-G. & Gilbert, C. Horizontal transfer and evolution of transposable elements in vertebrates. Nat. Commun. 11, 1362 (2020).

91. Charif, D. & Lobry, J. R. SeqinR 1.0-2: A Contributed Package to the R Project for Statistical Computing Devoted to Biological Sequences Retrieval and Analysis. in Structural Approaches to Sequence Evolution (eds Bastolla, U., Porto, M., Roman, H. E. & Vendruscolo, M.) 207–232 (Springer Berlin Heidelberg, Berlin, Heidelberg, 2007). doi:10.1007/978-3-540-35306-5_10.

92. Camacho, C. et al. BLAST+: architecture and applications. BMC Bioinformatics 10, 421 (2009).

93. Huerta-Cepas, J., Serra, F. & Bork, P. ETE 3: Reconstruction, Analysis, and Visualization of Phylogenomic Data. Mol. Biol. Evol. 33, 1635–1638 (2016).

