## Supplementary Figure for "Transposon-driven pervasive genome restructuring and regulatory evolution in early-diverging Holomycota"

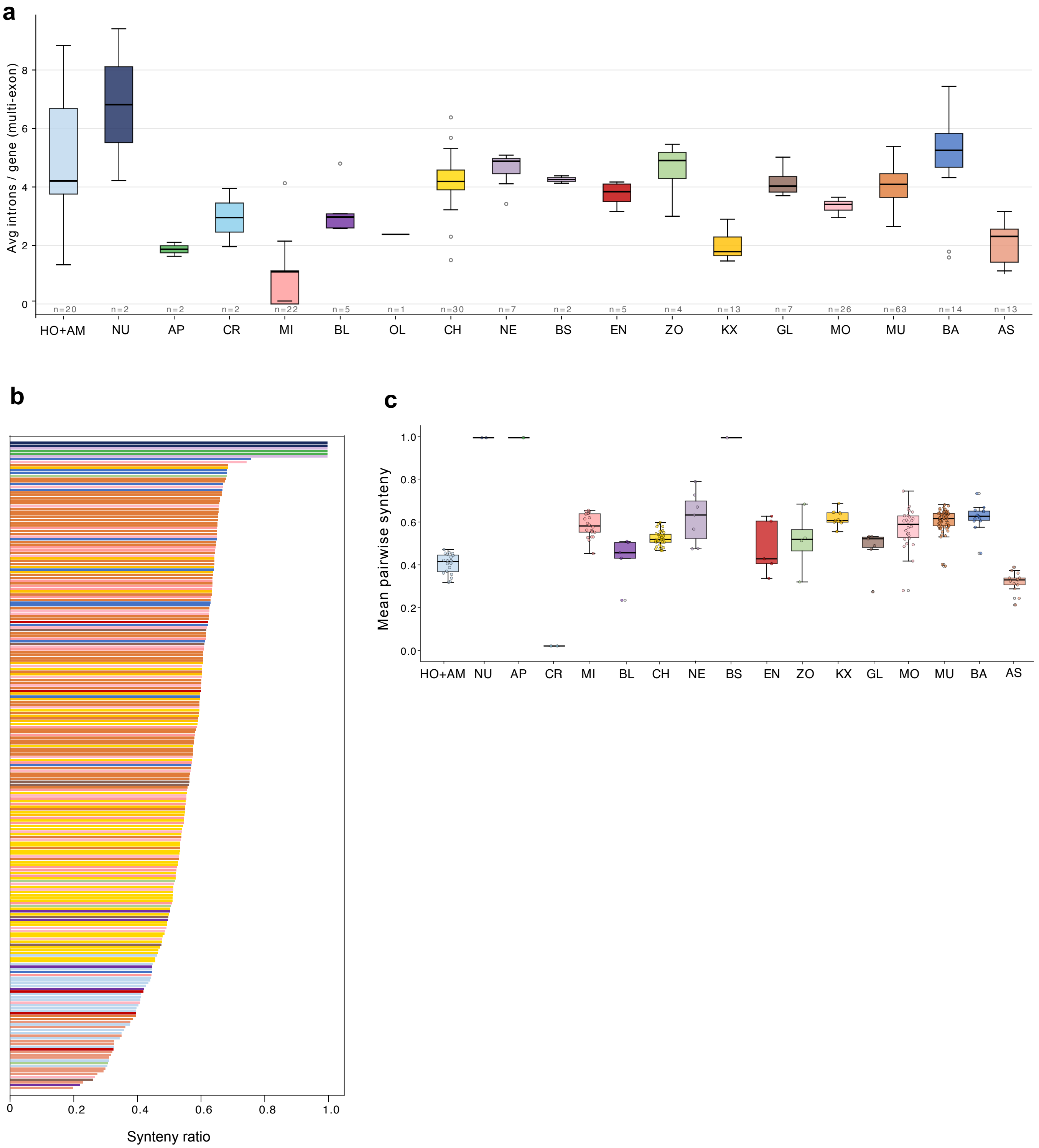

a
Avg introns / gene (multi-exon)
0
2
4
6
8
n=20
n=2
n=2
n=2
n=22
n=5
n=1
n=30
n=7
n=2
n=5
n=4
n=13
n=7
n=26
n=63
n=14
n=13
HO+AM
NU
AP
CR
MI
BL
OL
CH
NE
BS
EN
ZO
KX
GL
MO
MU
BA
AS
b
0
0.2
0.4
0.6
0.8
1.0
Synteny ratio
c
Mean pairwise synteny
0.0
0.2
0.4
0.6
0.8
1.0
HO+AM
NU
AP
CR
MI
BL
CH
NE
BS
EN
ZO
KX
GL
MO
MU
BA
AS

**Supplementary Figure 1.** (a) Box plot of mean introns per gene in multi-exon genes per clade across 18 NDF clades. Species counts shown below each box; box = IQR, whiskers = 1.5 × IQR. (b) Distribution of corrected pairwise synteny ratios (0–1.0) per species pair, bars coloured by comparison clade. (c) Box plot of mean pairwise corrected synteny ratio per clade (0.0–1.0).

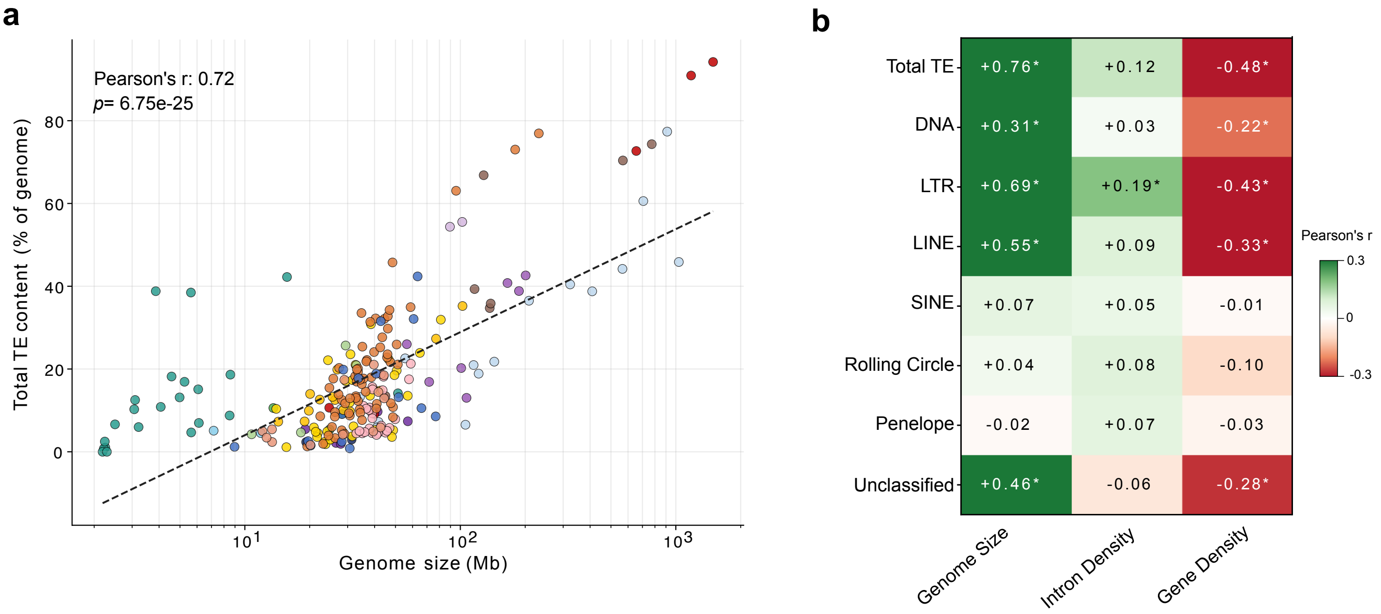
a
Pearson's r: 0.72
p= 6.75e-25
Total TE content (% of genome)
80
60
40
20
0
10^1
10^2
10^3
Genome size (Mb)
b
Total TE
DNA
LTR
LINE
SINE
Rolling Circle
Penelope
Unclassified
+0.76*
+0.12
-0.48*
+0.31*
+0.03
-0.22*
+0.69*
+0.19*
-0.43*
+0.55*
+0.09
-0.33*
+0.07
+0.05
-0.01
+0.04
+0.08
-0.10
-0.02
+0.07
-0.03
+0.46*
-0.06
-0.28*
Genome Size
Intron Density
Gene Density
Pearson's r
0.3
0
-0.3

**Supplementary Figure 2.** (a) Scatter plot of genome size (Mb, log scale, x-axis) versus total TE content (% of genome, y-axis). Points colored by clade; Pearson's r = 0.72, $p = 6.75 \times 10^{-25}$. (b) Heatmap of Pearson r values between eight TE category abundances and genomic properties. Asterisks = BH-corrected significance (FDR < 0.05)

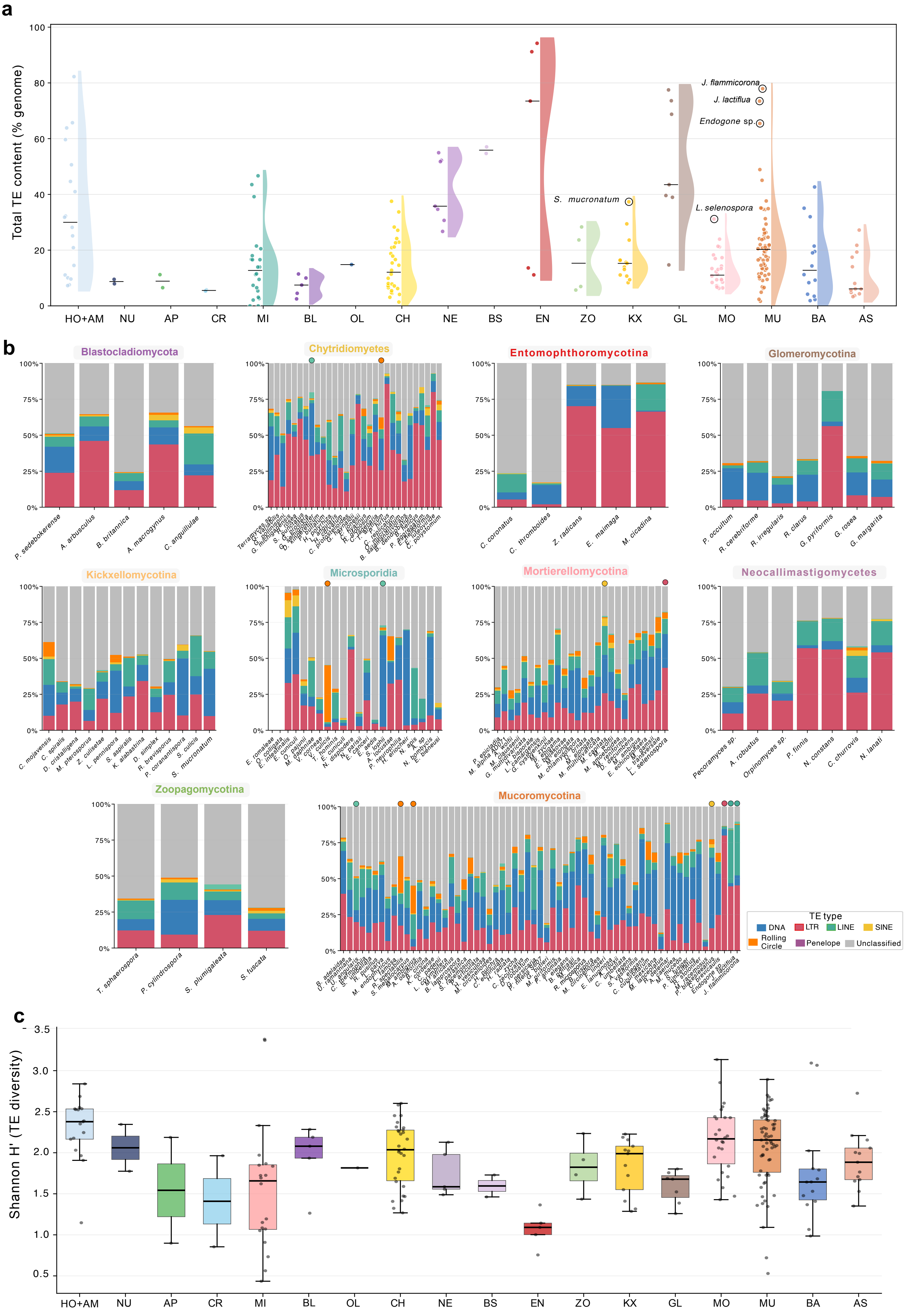
a
Total TE content (% genome)
S. mucronatum
J. flammicorona
J. lactiflua
Endogone sp.
L. selenospora
HO+AM
NU
AP
CR
MI
BL
OL
CH
NE
BS
EN
ZO
KX
GL
MO
MU
BA
AS
b
Blastocladiomycota
Chytridiomyetes
Entomophthoromycotina
Glomeromycotina
Kickxellomycotina
Microsporidia
Mortierellomycotina
Neocallimastigomycetes
Zoopagomycotina
Mucoromycotina
P. sedebokerense
A. arbusculus
B. britannica
A. macrogynus
C. anguillulae
C. coronatus
C. thromboides
Z. radicans
E. maimaga
M. cicadina
P. occultum
R. cerebriforme
R. irregularis
R. clarus
G. pyriformis
G. rosea
G. margarita
Pecoramyces sp.
A. robustus
Orpinomyces sp.
P. finnis
N. constans
C. churrovis
N. lanati
T. sphaerospora
P. cylindrospora
S. plumigaleata
S. fuscata
TE type
DNA
LTR
LINE
SINE
Rolling Circle
Penelope
Unclassified
c
Shannon H' (TE diversity)
HO+AM
NU
AP
CR
MI
BL
OL
CH
NE
BS
EN
ZO
KX
GL
MO
MU
BA
AS

**Supplementary Figure 3.** (a) Violin plots of total TE content (% genome) per clade. Outlier species in TE content for their clade are labeled. (b) Stacked bar charts of TE class composition per species. (c) Box plot of Shannon TE diversity index H′ per clade.

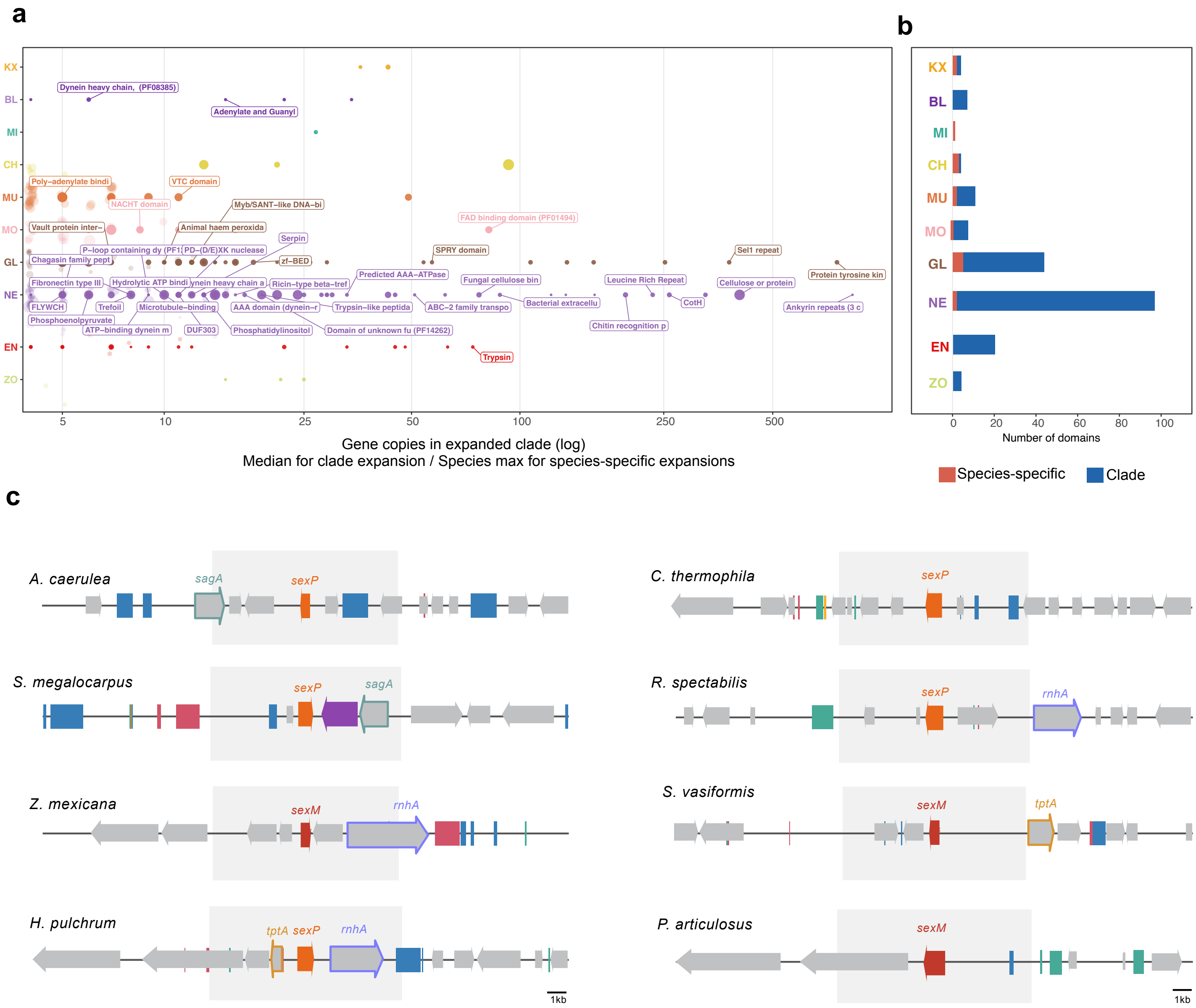
a
KX
BL
MI
CH
MU
MO
GL
NE
EN
ZO
Dynein heavy chain, (PF08385)
Adenylate and Guanyl
Poly−adenylate bindi
VTC domain
NACHT domain
Myb/SANT−like DNA−bi
FAD binding domain (PF01494)
Vault protein inter−
Animal haem peroxida
Serpin
P−loop containing dy (PF1
PD−(D/E)XK nuclease
SPRY domain
Sel1 repeat
Chagasin family pept
zf−BED
Protein tyrosine kin
Predicted AAA−ATPase
Fibronectin type III
Hydrolytic ATP bindi
ynein heavy chain a
Ricin−type beta−tref
Fungal cellulose bin
Leucine Rich Repeat
Cellulose or protein
FLYWCH
Trefoil
Microtubule−binding
AAA domain (dynein−r
Trypsin−like peptida
ABC−2 family transpo
Bacterial extracellu
CotH
Ankyrin repeats (3 c
Phosphoenolpyruvate
ATP−binding dynein m
DUF303
Phosphatidylinositol
Domain of unknown fu (PF14262)
Chitin recognition p
Trypsin
5
10
25
50
100
250
500
Gene copies in expanded clade (log)
Median for clade expansion / Species max for species-specific expansions
b
KX
BL
MI
CH
MU
MO
GL
NE
EN
ZO
0
20
40
60
80
100
Number of domains
Species-specific
Clade
c
A. caerulea
sagA
sexP
C. thermophila
sexP
S. megalocarpus
sexP
sagA
R. spectabilis
sexP
rnhA
Z. mexicana
sexM
rnhA
S. vasiformis
sexM
tptA
H. pulchrum
tptA
sexP
rnhA
P. articulosus
sexM
1kb
1kb

**Supplementary Figure 4.** (a) Domain-expanded gene families. TE-associated expansions are labelled (b) Count of expanded domains per clade, split by species-specific expansions (orange) and clade expansions (blue). (c) Mating type loci in MU species, genes and TEs are indicated for each species. Scale bar = 1 kb.

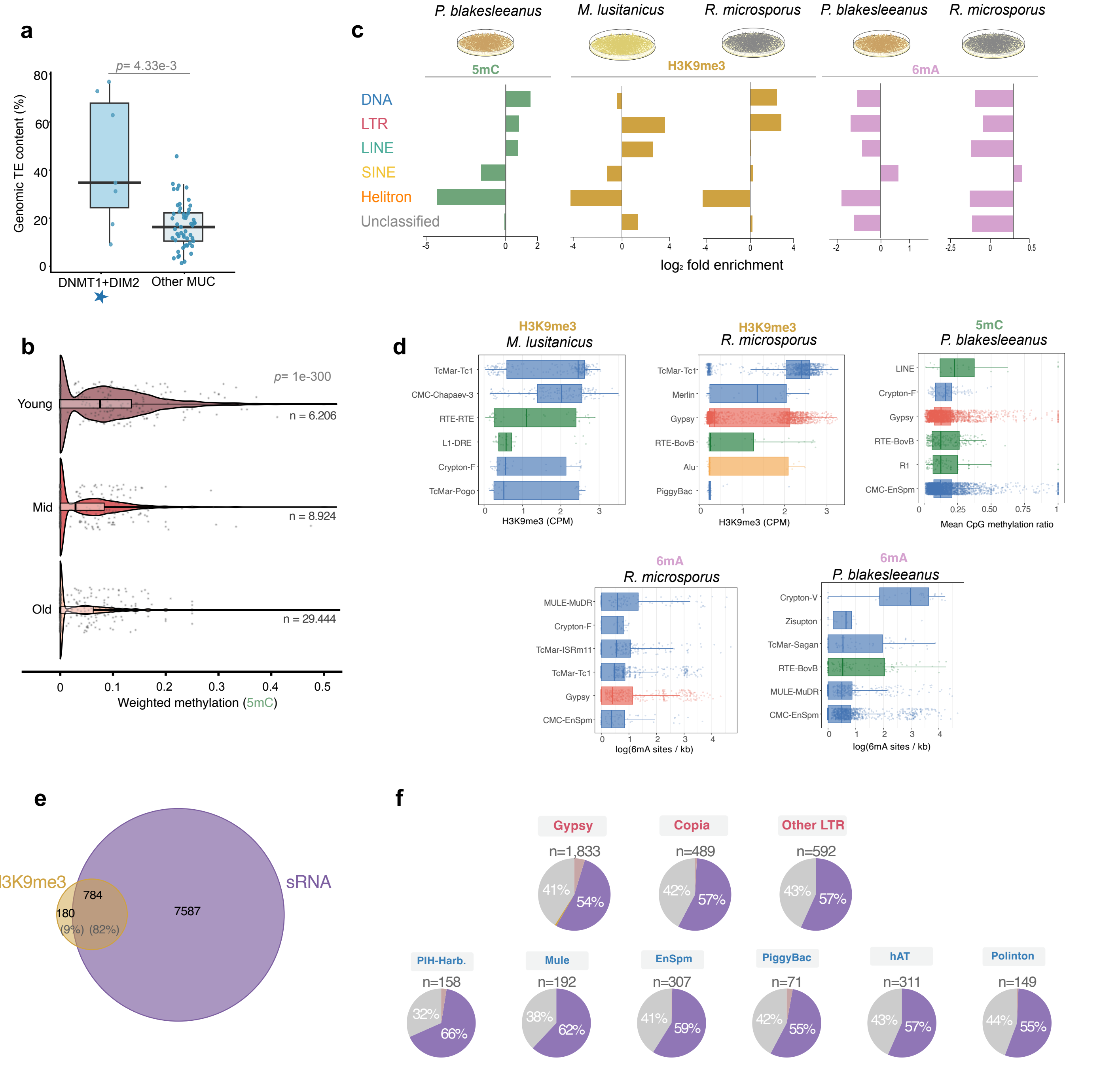
a
p= 4.33e-3
Genomic TE content (%)
0
20
40
60
80
DNMT1+DIM2
Other MUC
c
P. blakesleeanus
M. lusitanicus
R. microsporus
P. blakesleeanus
R. microsporus
5mC
H3K9me3
6mA
DNA
LTR
LINE
SINE
Helitron
Unclassified
log₂ fold enrichment
b
Young
Mid
Old
p= 1e-300
n = 6.206
n = 8.924
n = 29.444
0
0.1
0.2
0.3
0.4
0.5
Weighted methylation (5mC)
d
H3K9me3
M. lusitanicus
TcMar-Tc1
CMC-Chapaev-3
RTE-RTE
L1-DRE
Crypton-F
TcMar-Pogo
H3K9me3 (CPM)
H3K9me3
R. microsporus
TcMar-Tc1
Merlin
Gypsy
RTE-BovB
Alu
PiggyBac
H3K9me3 (CPM)
5mC
P. blakesleeanus
LINE
Crypton-F
Gypsy
RTE-BovB
R1
CMC-EnSpm
Mean CpG methylation ratio
6mA
R. microsporus
MULE-MuDR
Crypton-F
TcMar-ISRm11
TcMar-Tc1
Gypsy
CMC-EnSpm
log(6mA sites / kb)
6mA
P. blakesleeanus
Crypton-V
Zisupton
TcMar-Sagan
RTE-BovB
MULE-MuDR
CMC-EnSpm
log(6mA sites / kb)
e
H3K9me3
sRNA
784
180
7587
(9%)
(82%)
f
Gypsy
n=1,833
41%
54%
Copia
n=489
42%
57%
Other LTR
n=592
43%
57%
PIH-Harb.
n=158
32%
66%
Mule
n=192
38%
62%
EnSpm
n=307
41%
59%
PiggyBac
n=71
42%
55%
hAT
n=311
43%
57%
Polinton
n=149
44%
55%

**Supplementary Figure 5.** (a) Total TE genomic content (%) between Mucoromycota species carrying both DNMT1 and DIM2 orthologs versus all other Mucoromycota. Wilcoxon rank-sum test: $p = 4.33 \times 10^{-3}$. (b) Coverage-weighted 5mC methylation by TE age class: Young (n = 6,206 elements), Mid (n = 8,924), Old (n = 29,444). Kruskal–Wallis test: $p < 2.2 \times 10^{-300}$. (c) Enrichment (log2FC) in each epigenetic mark for each. (d) Per-TE-family box plots of epigenetic enrichment across different species and epigenetic modifications. (e) Euler diagram of TE loci enriched for H3K9me3 and targeted by sRNA. (f) Pie charts per TE family showing the proportion of TE copies targeted exclusively by sRNA versus co-targeted by both sRNA and H3K9me3.

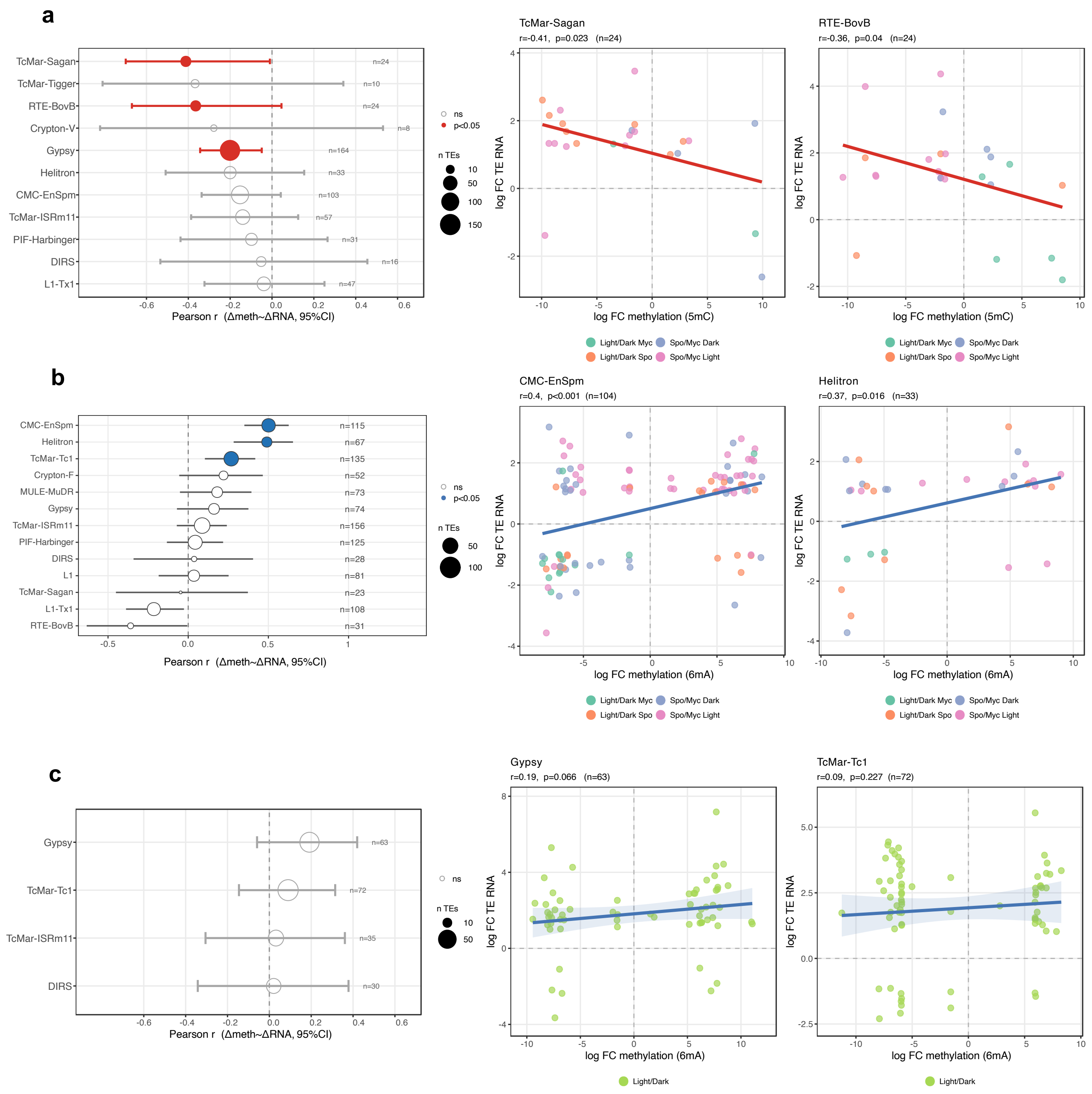

a
TcMar-Sagan
TcMar-Tigger
RTE-BovB
Crypton-V
Gypsy
Helitron
CMC-EnSpm
TcMar-ISRm11
PIF-Harbinger
DIRS
L1-Tx1
n=24
n=10
n=24
n=8
n=164
n=33
n=103
n=57
n=31
n=16
n=47
Pearson r (Δmeth~ΔRNA, 95%CI)
ns
p<0.05
n TEs
10
50
100
150
TcMar-Sagan
r=-0.41, p=0.023 (n=24)
RTE-BovB
r=-0.36, p=0.04 (n=24)
log FC TE RNA
log FC methylation (5mC)
Light/Dark Myc
Spo/Myc Dark
Light/Dark Spo
Spo/Myc Light
b
CMC-EnSpm
Helitron
TcMar-Tc1
Crypton-F
MULE-MuDR
Gypsy
TcMar-ISRm11
PIF-Harbinger
DIRS
L1
TcMar-Sagan
L1-Tx1
RTE-BovB
n=115
n=67
n=135
n=52
n=73
n=74
n=156
n=125
n=28
n=81
n=23
n=108
n=31
Pearson r (Δmeth~ΔRNA, 95%CI)
ns
p<0.05
n TEs
50
100
CMC-EnSpm
r=0.4, p<0.001 (n=104)
Helitron
r=0.37, p=0.016 (n=33)
log FC TE RNA
log FC methylation (6mA)
Light/Dark Myc
Spo/Myc Dark
Light/Dark Spo
Spo/Myc Light
c
Gypsy
TcMar-Tc1
TcMar-ISRm11
DIRS
n=63
n=72
n=35
n=30
Pearson r (Δmeth~ΔRNA, 95%CI)
ns
n TEs
10
50
Gypsy
r=0.19, p=0.066 (n=63)
TcMar-Tc1
r=0.09, p=0.227 (n=72)
log FC TE RNA
log FC methylation (6mA)
Light/Dark

**Supplementary Figure 6.** (a) Correlation (Pearson) of condition-matched changes in TE methylation (Δ5mC) and TE expression (ΔRNA) for 11 TE families; filled red circles = $p < 0.05$, TcMar-Sagan: $r = 0.41$, $p = 0.023$, $n = 24$; RTE-BovB: $r = -0.36$, $p = 0.04$, $n = 24$). (b) Correlation (Pearson) of condition-matched changes in TE methylation (Δ6mA) and TE expression (ΔRNA) 6mA context; filled blue circles = $p < 0.05$, TcMar-Tc1 ($r = 0.47$, one-tailed $p < 0.001$; $n = 135$), Helitron ($r = 0.47$, $p = 0.0002$; $n = 59$, CMC-EnSpm ($r = 0.38$, $p < 0.001$; $n= 115$). (c) Correlation (Pearson) of condition-matched changes in TE methylation (Δ6mA) and TE expression (ΔRNA) 6mA context for *R. microsporus*. Shown for reference: Gypsy $r = 0.19$, $p = 0.066$, $n = 63$; TcMar-Tc1 $r = -0.09$, $p = 0.227$, $n = 72$.

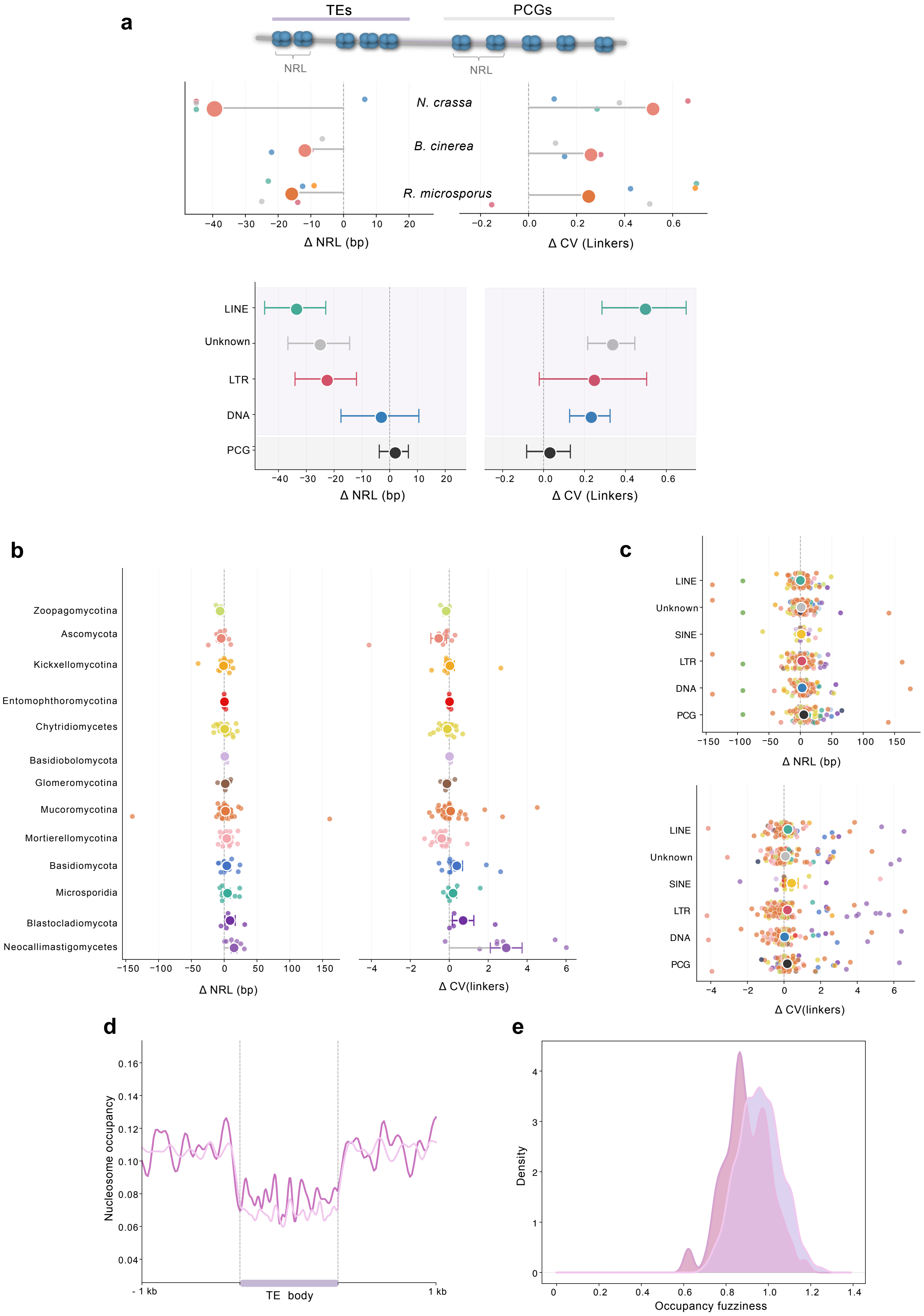
a
TEs
PCGs
NRL
NRL
N. crassa
B. cinerea
R. microsporus
Δ NRL (bp)
Δ CV (Linkers)
LINE
Unknown
LTR
DNA
PCG
Δ NRL (bp)
Δ CV (Linkers)
b
Zoopagomycotina
Ascomycota
Kickxellomycotina
Entomophthoromycotina
Chytridiomycetes
Basidiobolomycota
Glomeromycotina
Mucoromycotina
Mortierellomycotina
Basidiomycota
Microsporidia
Blastocladiomycota
Neocallimastigomycetes
Δ NRL (bp)
Δ CV(linkers)
c
LINE
Unknown
SINE
LTR
DNA
PCG
Δ NRL (bp)
LINE
Unknown
SINE
LTR
DNA
PCG
Δ CV(linkers)
d
Nucleosome occupancy
- 1 kb
TE body
1 kb
e
Density
Occupancy fuzziness

**Supplementary Figure 7.** (a) ΔNRL (bp) and ΔCV(linkers) with 95% confidence intervals for species with in vivo MNase-seq data (*Neurospora crassa*, *Botrytis cinerea*, *Rhizopus microsporus*), broken down by TE type (LINE, Unknown, LTR, DNA, PCG). (b) ΔNRL (bp) and ΔCV(linkers) for Nuclpred nucleosome occupancy predictions. Large filled dots = clade medians; small dots = individual species values. (c) Paired jitter plots of ΔNRL (bp) and ΔCV(linkers) broken down by TE type (large dots = clade medians). 95% CIs included. (d) Nucleosome occupancy profile of TEs with and without a MAC (±1 kb; body scaled). (e) Density distribution of profile fuzziness (1 − Pearson's r) of each TE relative to its group's mean profile. Higher values indicate greater heterogeneity in nucleosome positioning across TEs. Differences between groups are significant (Kolmogorov-Smirnov test $p < 0.001$).

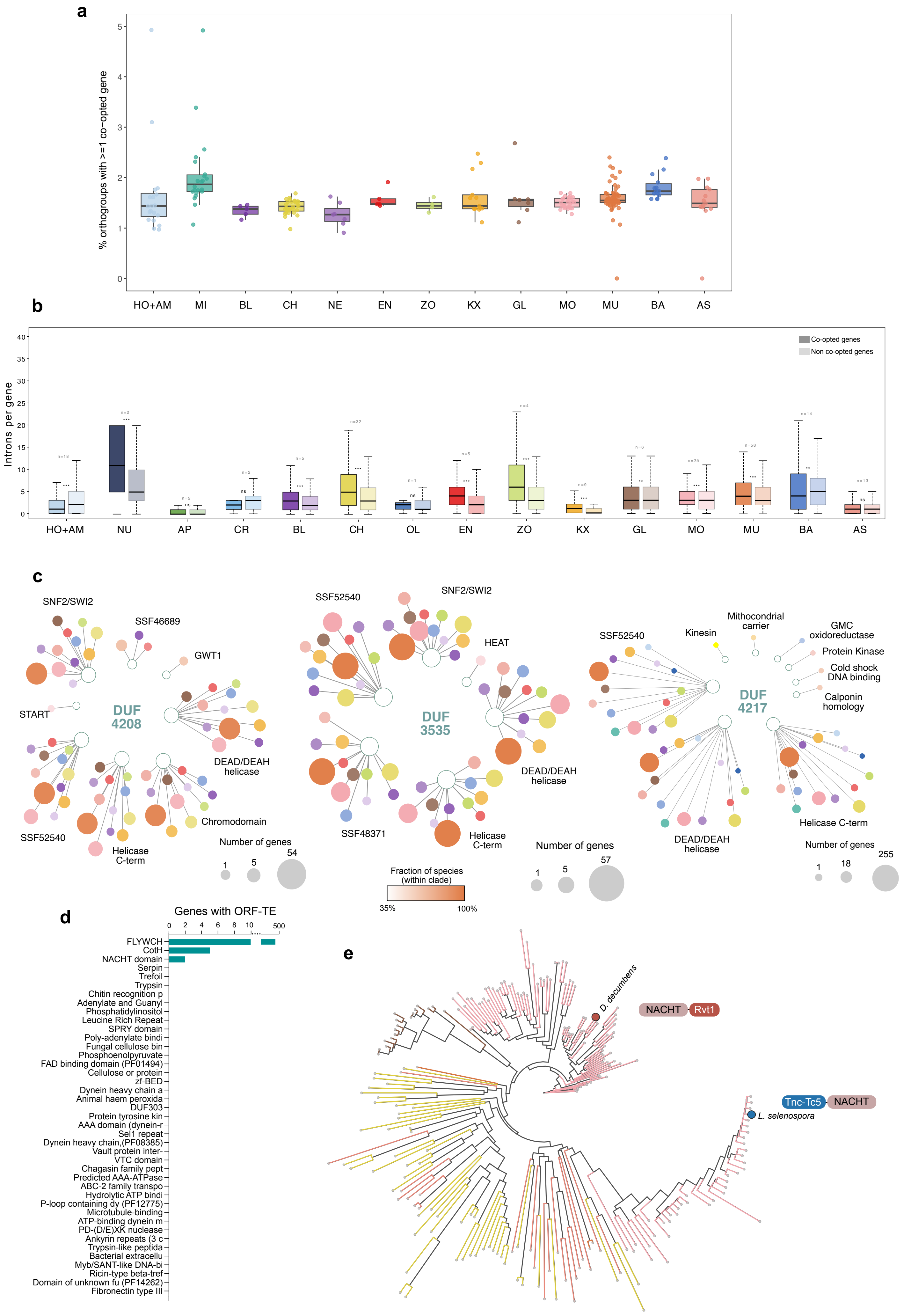
a
% orthogroups with >=1 co−opted gene
HO+AM
MI
BL
CH
NE
EN
ZO
KX
GL
MO
MU
BA
AS
b
Introns per gene
Co-opted genes
Non co-opted genes
HO+AM
NU
AP
CR
BL
CH
OL
EN
ZO
KX
GL
MO
MU
BA
AS
c
SNF2/SWI2
SSF46689
GWT1
START
DUF 4208
SSF52540
Helicase C-term
Chromodomain
DEAD/DEAH helicase
Number of genes
SSF52540
SNF2/SWI2
HEAT
DUF 3535
SSF48371
Helicase C-term
DEAD/DEAH helicase
Fraction of species (within clade)
35%
100%
Number of genes
SSF52540
Kinesin
Mithocondrial carrier
GMC oxidoreductase
Protein Kinase
Cold shock DNA binding
Calponin homology
DUF 4217
DEAD/DEAH helicase
Helicase C-term
Number of genes
d
Genes with ORF-TE
FLYWCH
CotH
NACHT domain
Serpin
Trefoil
Trypsin
Chitin recognition p
Adenylate and Guanyl
Phosphatidylinositol
Leucine Rich Repeat
SPRY domain
Poly-adenylate bindi
Fungal cellulose bin
Phosphoenolpyruvate
FAD binding domain (PF01494)
Cellulose or protein
zf-BED
Dynein heavy chain a
Animal haem peroxida
DUF303
Protein tyrosine kin
AAA domain (dynein-r
Sel1 repeat
Dynein heavy chain,(PF08385)
Vault protein inter-
VTC domain
Chagasin family pept
Predicted AAA-ATPase
ABC-2 family transpo
Hydrolytic ATP bindi
P-loop containing dy (PF12775)
Microtubule-binding
ATP-binding dynein m
PD-(D/E)XK nuclease
Ankyrin repeats (3 c
Trypsin-like peptida
Bacterial extracellu
Myb/SANT-like DNA-bi
Ricin-type beta-tref
Domain of unknown fu (PF14262)
Fibronectin type III
e
D. decumbens
NACHT
Rvt1
Tnc-Tc5
NACHT
L. selenospora

**Supplementary Figure 8.** (a) Percentage of orthogroups containing ≥1 co-opted gene per clade. (b) Paired box plots of introns per gene for co-opted (darker) versus non-co-opted (lighter) genes per clade. Significance: ** $p < 0.01$, *** $p < 0.001$ (Wilcoxon signed-rank test, BH correction). (c) Domain interaction network diagrams for DUF4208, DUF3535, and DUF4217. (d) Number of genes carrying each ORF-derived TE domain fusion across NDF species. (e) Phylogenetic tree annotated with fusion events of the NACHT domain.

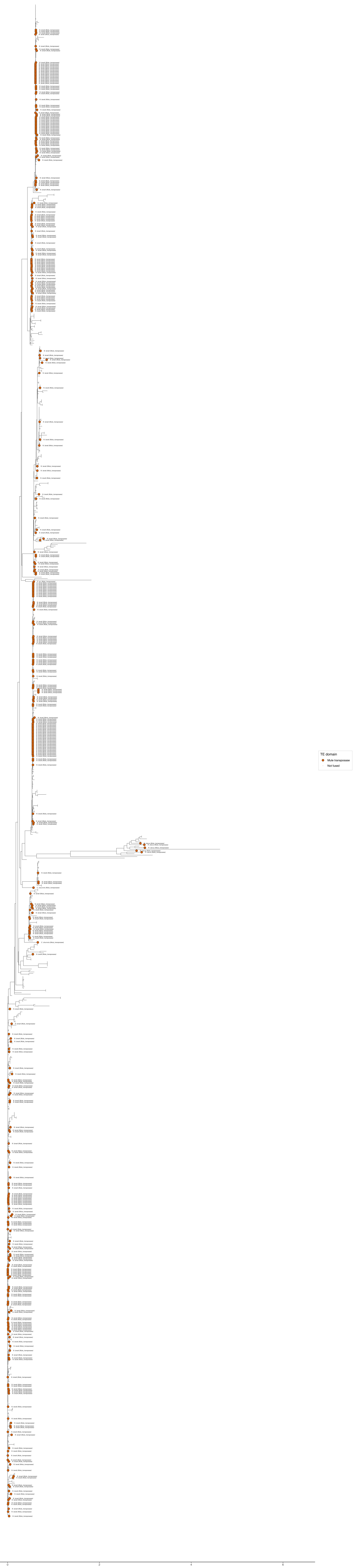

TE domain
Mule transposase
Not fused
0
2
4
6

**Supplementary Figure 9.** (a) Full phylogenetic tree of FLYWCH domains. Co-opted genes are indicated with orange dots.

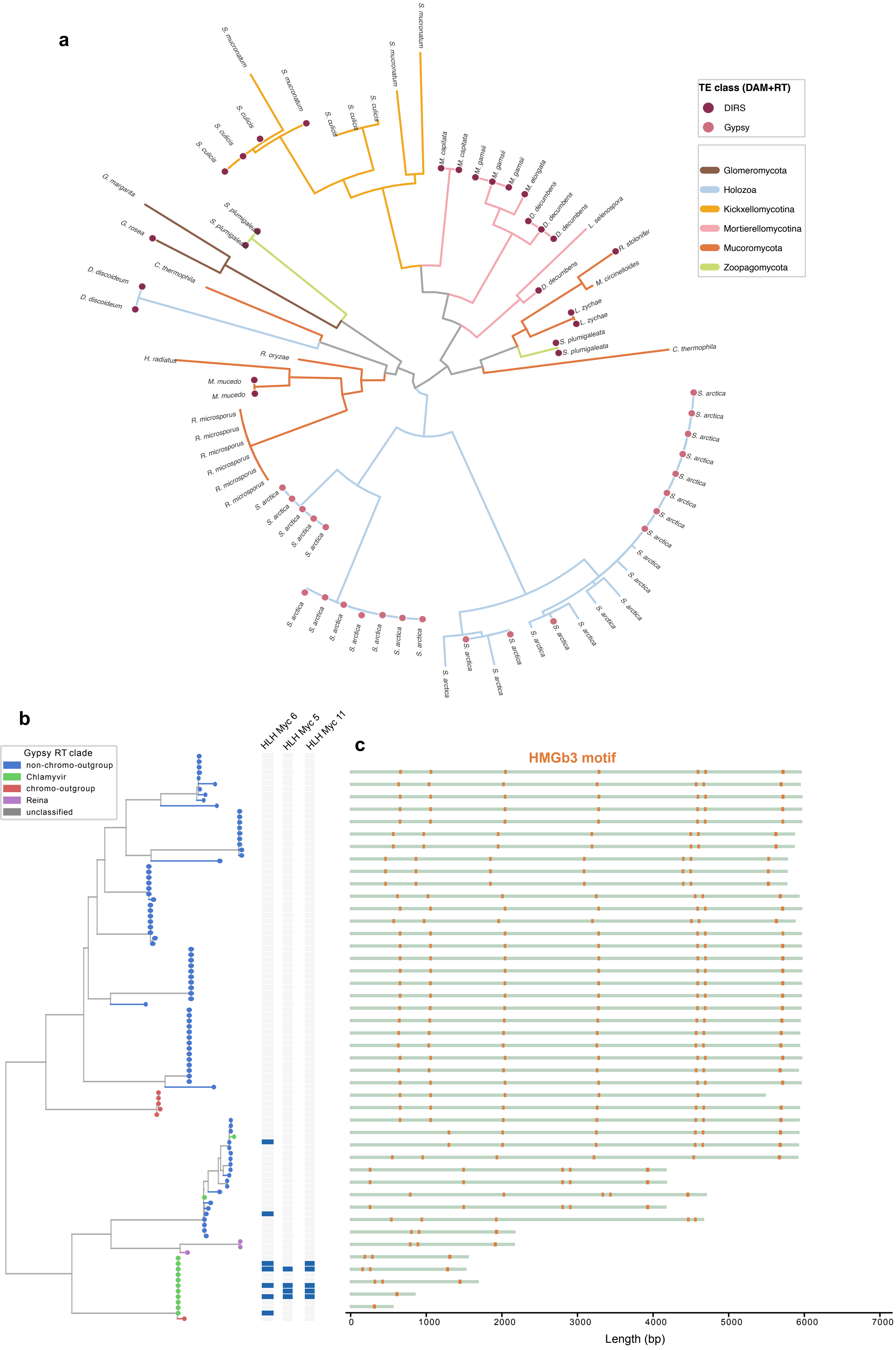
a
TE class (DAM+RT)
DIRS
Gypsy
Glomeromycota
Holozoa
Kickxellomycotina
Mortierellomycotina
Mucoromycota
Zoopagomycota
S. mucronatum
S. culicis
S. plumigaleata
G. margarita
G. rosea
C. thermophila
D. discoideum
H. radiatus
R. oryzae
M. mucedo
R. microsporus
S. arctica
M. capitata
M. gamsii
M. elongata
D. decumbens
L. selenospora
R. stolonifer
M. circinelloides
L. zychae
S. plumigaleata
b
Gypsy RT clade
non-chromo-outgroup
Chlamyvir
chromo-outgroup
Reina
unclassified
HLH Myc 6
HLH Myc 5
HLH Myc 11
c
HMGb3 motif
0
1000
2000
3000
4000
5000
6000
7000
Length (bp)

**Supplementary Figure 10.** (a) Maximum-likelihood tree of all DAM domains identified in the dataset. Branches are colored by clade and tips circled by the type of TE involved in the co-option (DIRS/Gypsy). (b) Tree using RT domains for Gypsy copies in *R. microsporus*. Squares determine presence/absence of peaks overlapping each TE. (c) GremLINE elements identified in the *M. lusitanicus* pericentromeric regions. Orange regions mark FIMO-identified HMGb3 binding sites.

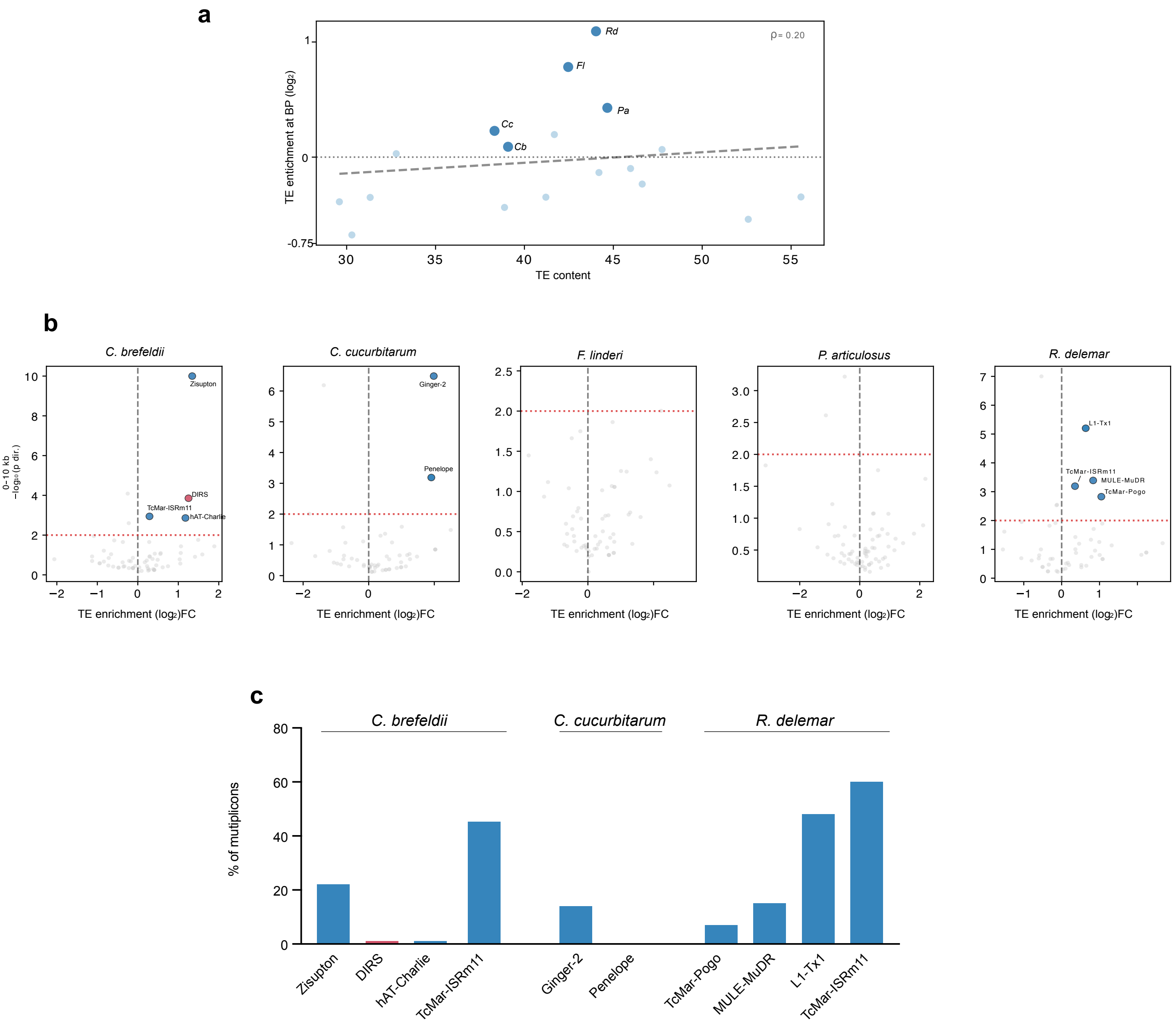

a
ρ= 0.20
Rd
Fl
Pa
Cc
Cb
TE enrichment at BP (log2)
TE content
b
C. brefeldii
C. cucurbitarum
F. linderi
P. articulosus
R. delemar
0–10 kb
−log10(p dir.)
Zisupton
DIRS
TcMar-ISRm11
hAT-Charlie
Ginger-2
Penelope
L1-Tx1
MULE-MuDR
TcMar-Pogo
TE enrichment (log2)FC
c
C. brefeldii
C. cucurbitarum
R. delemar
% of mutiplicons
Zisupton
DIRS
hAT-Charlie
TcMar-ISRm11
Ginger-2
Penelope
TcMar-Pogo
MULE-MuDR
L1-Tx1
TcMar-ISRm11

**Supplementary Figure 11.** (a) Scatter plot of total TE content versus TE enrichment at WGD synteny breakpoints per WGD species. Spearman $\rho = 20$, $p = 0.438$. (b) TE superfamily enrichment at synteny breakpoints and 10 kb flanking regions. (c) Percentage of syntenic multiplicons containing each significant TE superfamily in both copies of the duplicated segment.

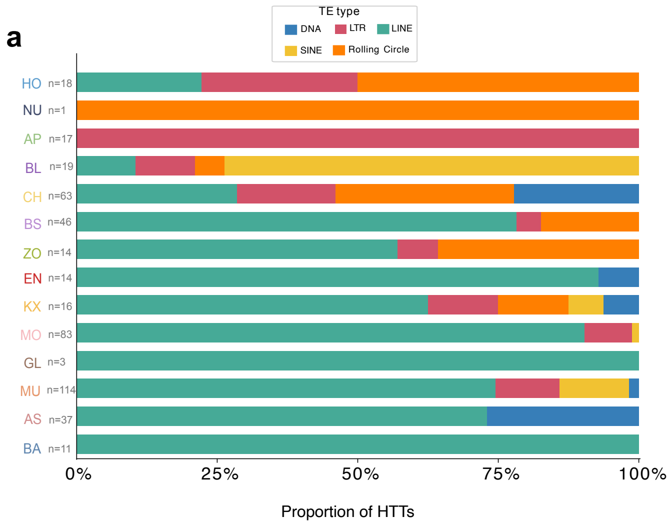

a
TE type
DNA
LTR
LINE
SINE
Rolling Circle
HO n=18
NU n=1
AP n=17
BL n=19
CH n=63
BS n=46
ZO n=14
EN n=14
KX n=16
MO n=83
GL n=3
MU n=114
AS n=37
BA n=11
0%
25%
50%
75%
100%
Proportion of HTTs
b
LINE
RC
LTR
SINE
DNA
Fraction of HTTs (%)
Genomic abudance (% of all TE)
0
10
20
30
40
50
60
70

**Supplementary Figure 12.** (a) Stacked bar chart of the proportion of HTT events by TE class. (b) Scatter plot of genomic TE class abundance (% of all TE) versus fraction of HTT events (% of all HTT events). Dashed diagonal = null expectation.

**Supplementary Table 1**: Domain co-opted in ORF fusion with TE domains, the number of occurrences and full domain composition of co-opted genes are indicated.

| Non-TE domain | TE domain | TE type | N# | Clade:Species | Domain architecture |
|---|---|---|---|---|---|
| FLYWCH | Mule_transposase | DNA transposon (MULE) | 385 | CH_Caecom1; CH_Neolan1; CH_Pecora1; GL_Rhicl1 | FLYWCH zinc finger domain \| MULE transposase domain |
| NACHT | HTH_Tnp_Tc5 | DNA transposon (hAT/Tc5) | 1 | MO_Morsel1 | Transposase \| NACHT domain \| HEAT repeats |
| NACHT | RVT_1 | LTR retrotransposon | 1 | MO_Disdec1 | Reverse transcriptase (RNA-dependent DNA polymerase) \| WD domain, G-beta repeat \| NACHT domain \| Heterokaryon |
| CotH | Mule_transposase | DNA transposon (MULE) | 2 | NE_Neolan1 | CotH protein \| MULE transposase domain |
| CotH | RVT_1 | LTR retrotransposon | 2 | NE_Orpsp1 | RVT_1 \| CotH |
| CotH | RVT_2 | LTR retrotransposon | 1 | CH_Orpsp1 | rve \| CBM_10 \| RVT_2 \| CotH |
| CotH | rve | LTR retrotransposon | 1 | CH_Orpsp1 | rve \| CBM_10 \| RVT_2 \| CotH |
