## Supplementary material for "Transposon-driven pervasive genome restructuring and regulatory evolution in early-diverging Holomycota": Supplementary File 2.pdf

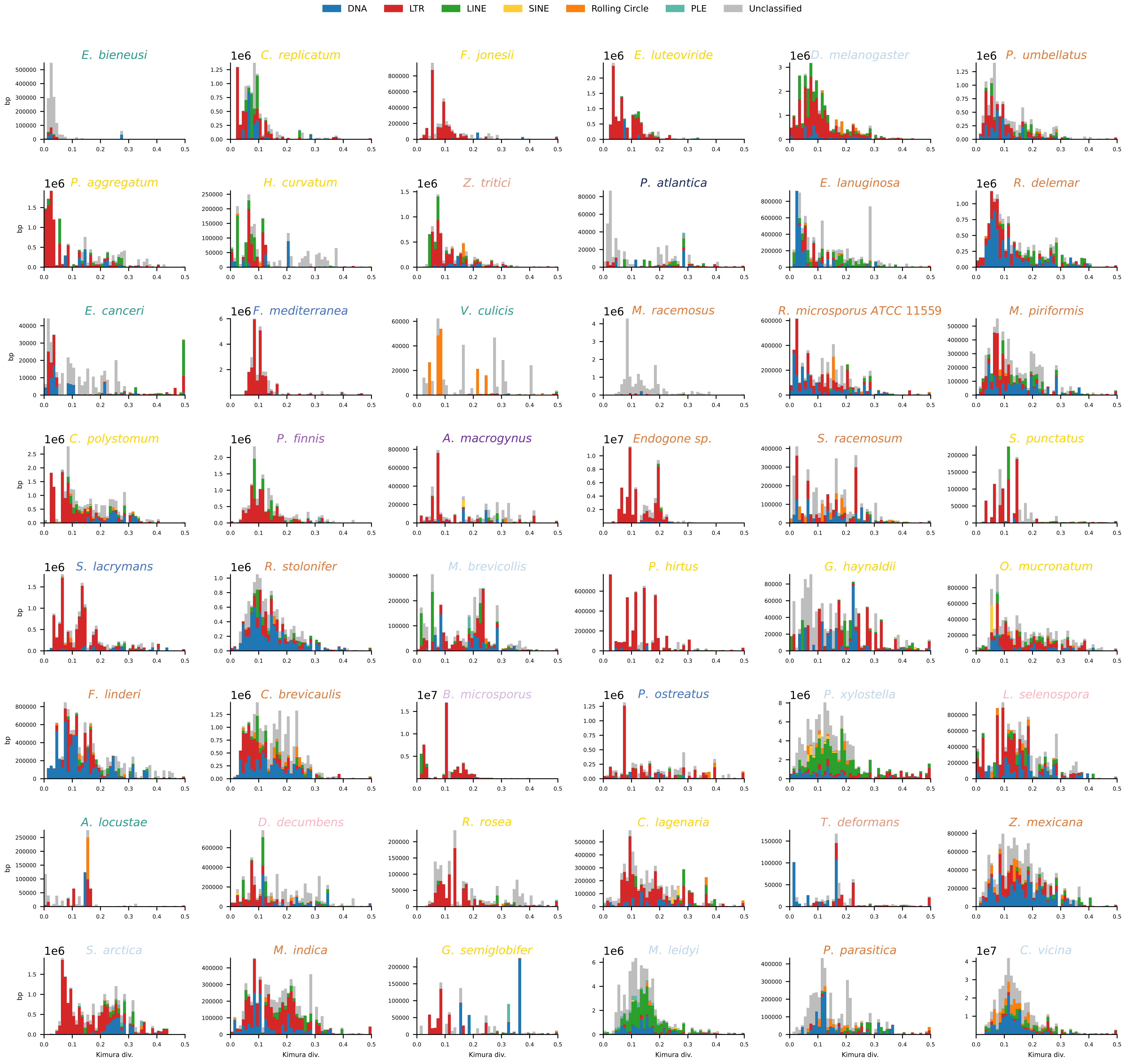
DNA
LTR
LINE
SINE
Rolling Circle
PLE
Unclassified
E. bieneusi
C. replicatum
F. jonesii
E. luteoviride
D. melanogaster
P. umbellatus
P. aggregatum
H. curvatum
Z. tritici
P. atlantica
E. lanuginosa
R. delemar
E. canceri
F. mediterranea
V. culicis
M. racemosus
R. microsporus ATCC 11559
M. piriformis
C. polystomum
P. finnis
A. macrogynus
Endogone sp.
S. racemosum
S. punctatus
S. lacrymans
R. stolonifer
M. brevicollis
P. hirtus
G. haynaldii
O. mucronatum
F. linderi
C. brevicaulis
B. microsporus
P. ostreatus
P. xylostella
L. selenospora
A. locustae
D. decumbens
R. rosea
C. lagenaria
T. deformans
Z. mexicana
S. arctica
M. indica
G. semiglobifer
M. leidyi
P. parasitica
C. vicina
bp
Kimura div.

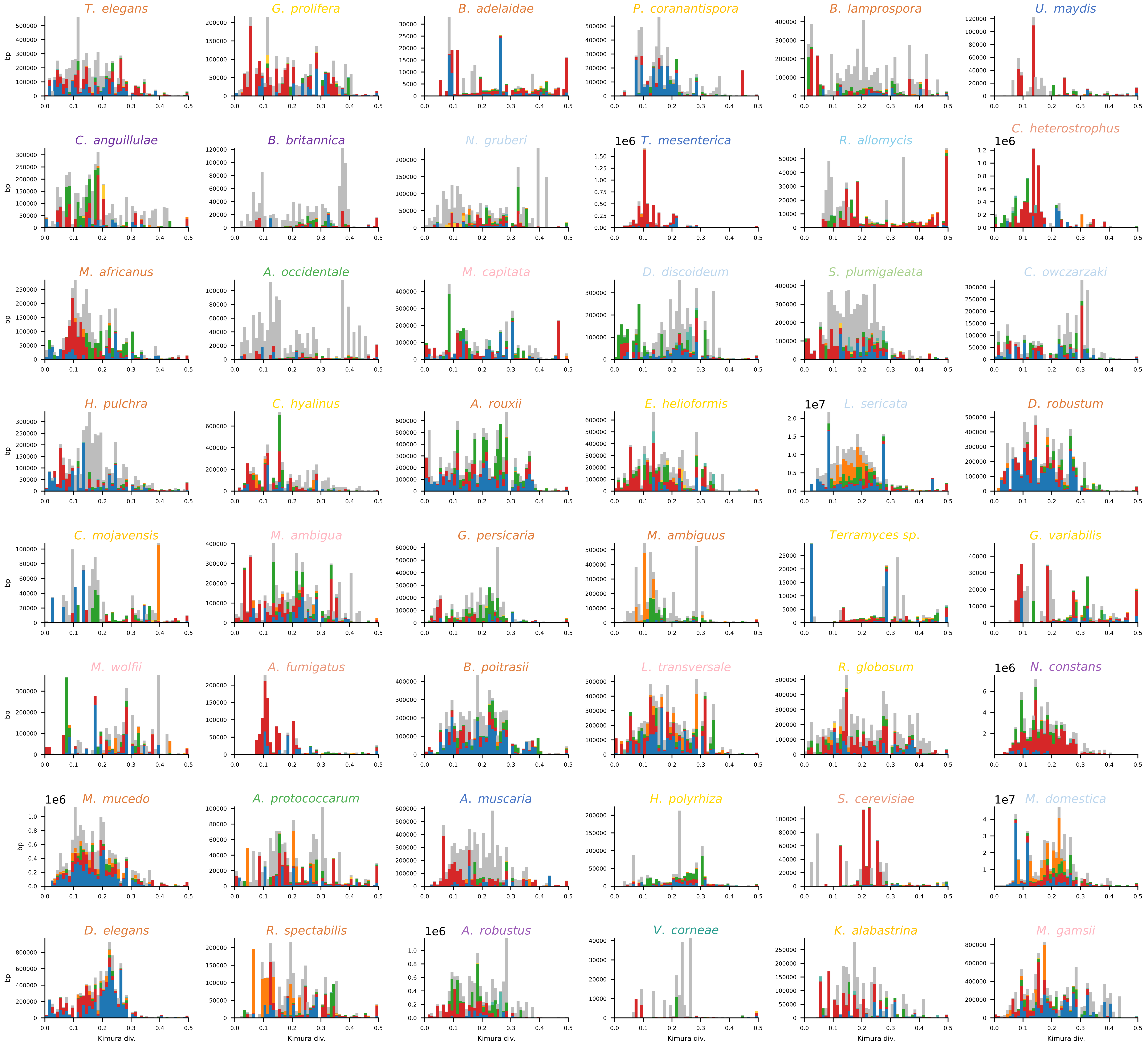
T. elegans
G. prolifera
B. adelaidae
P. coranantispora
B. lamprospora
U. maydis
C. anguillulae
B. britannica
N. gruberi
1e6 T. mesenterica
R. allomycis
C. heterostrophus
1e6
M. africanus
A. occidentale
M. capitata
D. discoideum
S. plumigaleata
C. owczarzaki
H. pulchra
C. hyalinus
A. rouxii
E. helioformis
1e7 L. sericata
D. robustum
C. mojavensis
M. ambigua
G. persicaria
M. ambiguus
Terramyces sp.
G. variabilis
M. wolfii
A. fumigatus
B. poitrasii
L. transversale
R. globosum
1e6 N. constans
1e6 M. mucedo
A. protococcarum
A. muscaria
H. polyrhiza
S. cerevisiae
1e7 M. domestica
D. elegans
R. spectabilis
1e6 A. robustus
V. corneae
K. alabastrina
M. gamsii
bp
Kimura div.

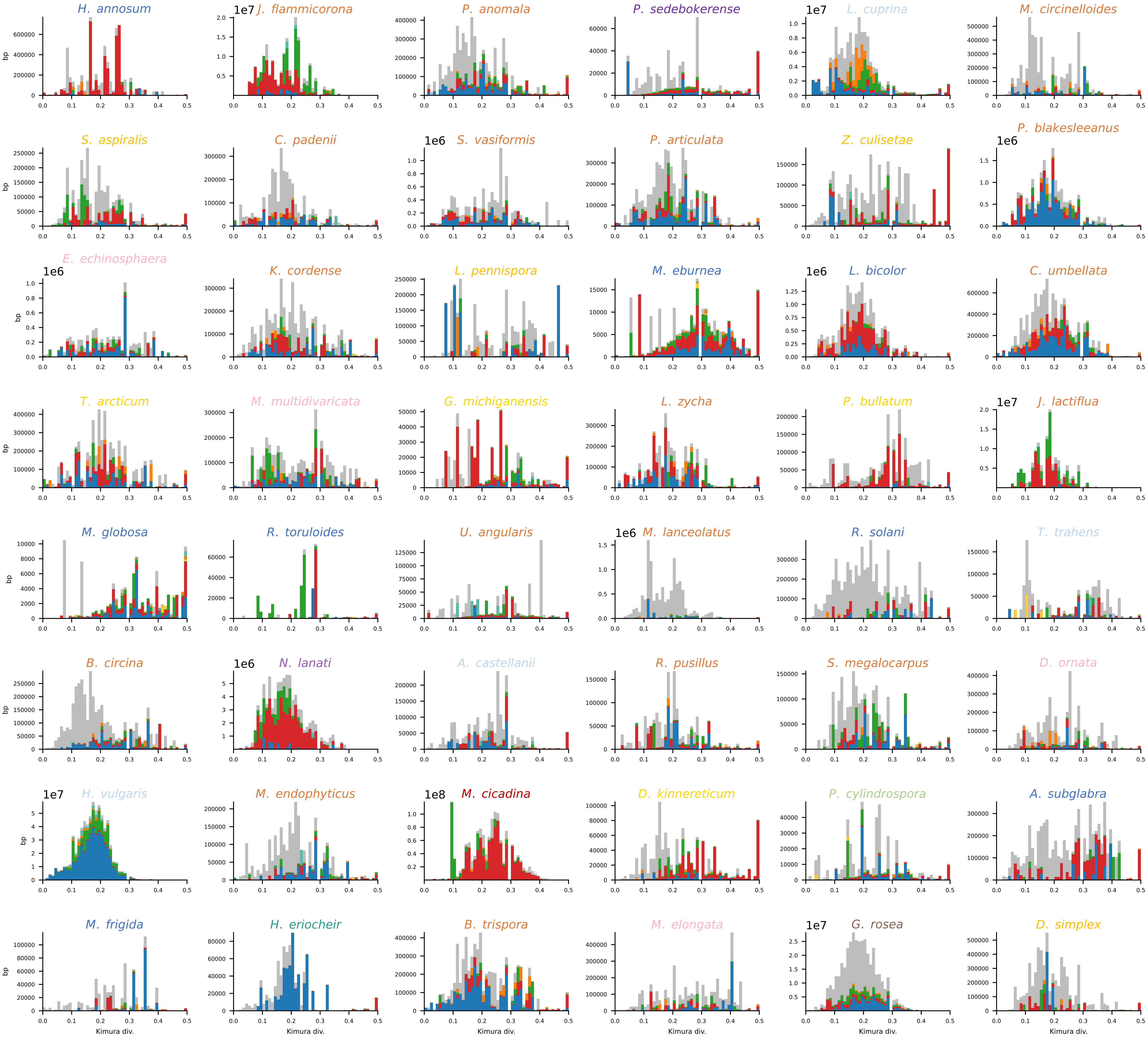

H. annosum
J. flammicorona
P. anomala
P. sedebokerense
L. cuprina
M. circinelloides
S. aspiralis
C. padenii
S. vasiformis
P. articulata
Z. culisetae
P. blakesleeanus
E. echinosphaera
K. cordense
L. pennispora
M. eburnea
L. bicolor
C. umbellata
T. arcticum
M. multidivaricata
G. michiganensis
L. zycha
P. bullatum
J. lactiflua
M. globosa
R. toruloides
U. angularis
M. lanceolatus
R. solani
T. trahens
B. circina
N. lanati
A. castellanii
R. pusillus
S. megalocarpus
D. ornata
H. vulgaris
M. endophyticus
M. cicadina
D. kinnereticum
P. cylindrospora
A. subglabra
M. frigida
H. eriocheir
B. trispora
M. elongata
G. rosea
D. simplex
bp
Kimura div.

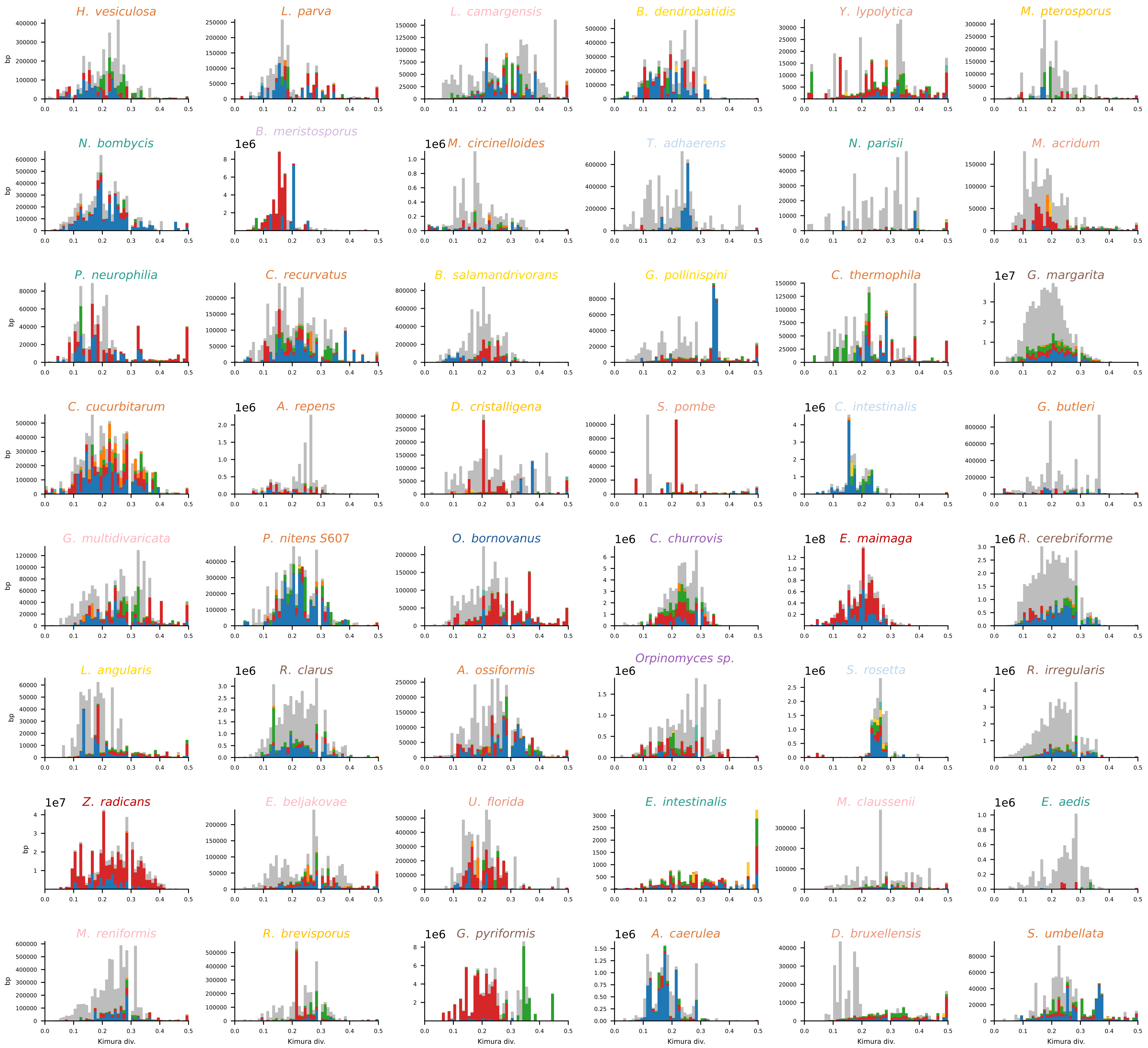

H. vesiculosa
L. parva
L. camargensis
B. dendrobatidis
Y. lypolytica
M. pterosporus
N. bombycis
B. meristosporus
M. circinelloides
T. adhaerens
N. parisii
M. acridum
P. neurophilia
C. recurvatus
B. salamandrivorans
G. pollinispini
C. thermophila
G. margarita
C. cucurbitarum
A. repens
D. cristalligena
S. pombe
C. intestinalis
G. butleri
G. multidivaricata
P. nitens S607
O. bornovanus
C. churrovis
E. maimaga
R. cerebriforme
L. angularis
R. clarus
A. ossiformis
Orpinomyces sp.
S. rosetta
R. irregularis
Z. radicans
E. beljakovae
U. florida
E. intestinalis
M. claussenii
E. aedis
M. reniformis
R. brevisporus
G. pyriformis
A. caerulea
D. bruxellensis
S. umbellata
bp
Kimura div.

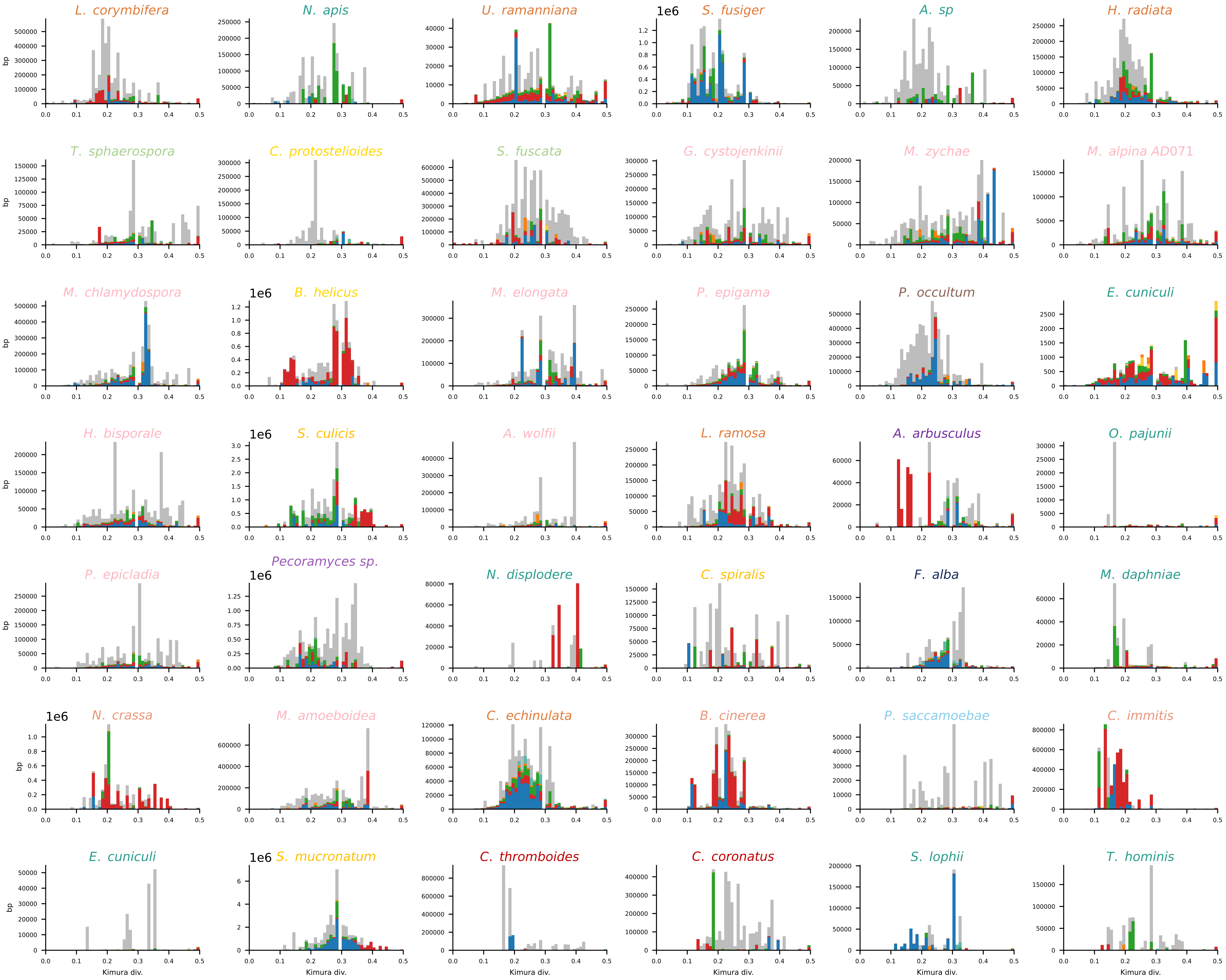
L. corymbifera
N. apis
U. ramanniana
S. fusiger
A. sp
H. radiata
T. sphaerospora
C. protostelioides
S. fuscata
G. cystojenkinii
M. zychae
M. alpina AD071
M. chlamydospora
B. helicus
M. elongata
P. epigama
P. occultum
E. cuniculi
H. bisporale
S. culicis
A. wolfii
L. ramosa
A. arbusculus
O. pajunii
P. epicladia
Pecoramyces sp.
N. displodere
C. spiralis
F. alba
M. daphniae
N. crassa
M. amoeboidea
C. echinulata
B. cinerea
P. saccamoebae
C. immitis
E. cuniculi
S. mucronatum
C. thromboides
C. coronatus
S. lophii
T. hominis
bp
1e6
Kimura div.
