## Supplementary material for "Transposon-driven pervasive genome restructuring and regulatory evolution in early-diverging Holomycota": Supplementary File 8.pdf

### Licensed Silhouettes

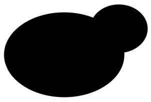

License: CC0 1.0 Universal Public Domain Dedication
Uploaded: 2013 Nov 10 by Wayne Decatur
Taxon: Saccharomyces cerevisiae Meyen ex E. C. Hansen 1883
Link: https://www.phylopic.org/images/347bc4ab-c1e0-4313-9ea1-939c2ad71232/saccharomyces-cerevisiae

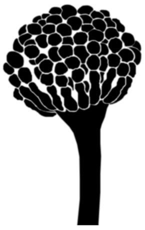

License: CC0 1.0 Universal Public Domain Dedication
Uploaded: 2019 Jun 16 by Mason McNai
Taxon: Aspergillus nidulans
Link: https://www.phylopic.org/images/c98aba13-4ba8-4448-bda1cf3500264954/aspergillus-nidulans

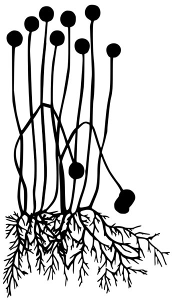

License: CC0 1.0 Universal Public Domain Dedication
Uploaded: 2023 Mar 31 by Guillaume Dera
Taxon: Mucor mucedo
Link: https://www.phylopic.org/images/ba2078ef-1e34-4181-a0a0-893abac714ad/mucor-mucedo

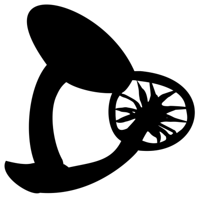

License: CC0 1.0 Universal Public Domain Dedication
Uploaded: 2022 Mar 25 by Riccardo percudani
Taxon: Laccaria Berkeley et Broome 1883
Link: https://www.phylopic.org/images/0ec7e6e6-e115-450a-b346-c40685ed3b1a/laccaria

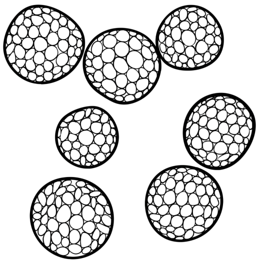

License: CC0 1.0 Universal Public Domain Dedication
Uploaded: 2025 Sep 30 by Arcadia Science
Taxon: Mycosarcoma maydis (DC.) Bref.
Link: https://www.phylopic.org/images/c2fad2f3-fa5a-496e-8388-2d71de3ff1e5/mycosarcoma-maydis

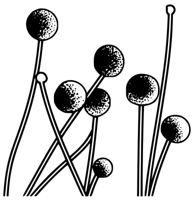

License: CC0 1.0 Universal Public Domain Dedication
Uploaded: 2023 Jan 15 by Levi Simons
Taxon: Umbelopsidomycetes
Link: https://www.phylopic.org/images/cb06a1f3-50f3-48fe-9797-c078bc040b4c/umbelopsidomycetes

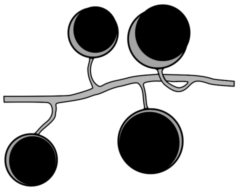

License: CC0 1.0 Universal Public Domain Dedication
Uploaded: 2015 Sep 20 by Yan Wong
Taxon: Glomus diaphanum
Link: https://www.phylopic.org/images/31aa0781-ca28-4362-8955-520f1c45f232/glomus-diaphanum

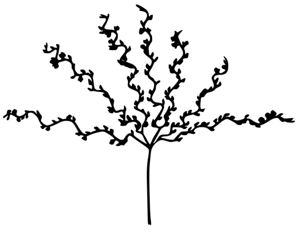

License: CC0 1.0 Universal Public Domain Dedication
Uploaded: 2023 Mar 31 by Guillaume Dera
Taxon: Coemansia umbellata
Link: https://www.phylopic.org/images/309c7adf-c2da-40fd-904b-e231c2d82dac/coemansia-umbellata

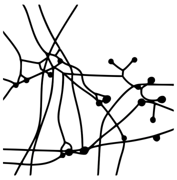

License: CC0 1.0 Universal Public Domain Dedication
Uploaded: 2023 Jan 13 by Levi Simons
Taxon: Zoopagales
Link: [https://www.phylopic.org/images/93318b1b-a277-48dd-b9c3-da2ef9d97eb9/zoopagales](https://www.phylopic.org/images/93318b1b-a277-48dd-b9c3-da2ef9d97eb9/zoopagales)

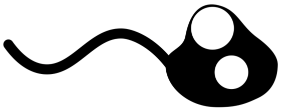

License: CC0 1.0 Universal Public Domain Dedication
Uploaded: 2022 Nov 8 by Andrew JM Swafford
Taxon: Spizellomyces
Link: [https://www.phylopic.org/images/15860a9d-0013-4214-b945-cfc96aba8266/spizellomyces](https://www.phylopic.org/images/15860a9d-0013-4214-b945-cfc96aba8266/spizellomyces)

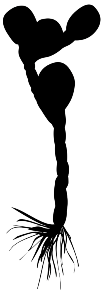

License: CC0 1.0 Universal Public Domain Dedication
Uploaded: 2012 Feb 29 by T. Michael Keesey
Taxon: Allomyces arbuscula E. J. Butler
Link: [https://www.phylopic.org/images/f72ec90f-d008-4908-a7e6-582e04814f95/allomyces-arbuscula](https://www.phylopic.org/images/f72ec90f-d008-4908-a7e6-582e04814f95/allomyces-arbuscula)

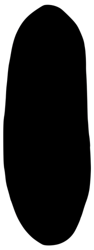

License: CC0 1.0 Universal Public Domain Dedication
Uploaded: 2023 Mar 31 by Guillaume Dera
Taxon: Nosema lepocreadii
Link: [https://www.phylopic.org/images/05def04a-f550-4b77-bdbe-d7c25a22f229/nosema-lepocreadii](https://www.phylopic.org/images/05def04a-f550-4b77-bdbe-d7c25a22f229/nosema-lepocreadii)

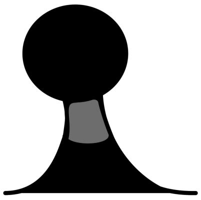

License: CC0 1.0 Universal Public Domain Dedication
Uploaded: 2015 Nov 23 by Yan Wong
Taxon: Fonticula alba
Link: https://www.phylopic.org/images/baaab011-0673-4d18-804a-f51394d29235/fonticula-alba

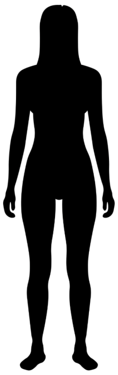

License: CC0 1.0 Universal Public Domain Dedication
Uploaded: 2023 Mar 15 by Carlo De Rito
Taxon: Homo sapiens sapiens
Link: https://www.phylopic.org/images/37536a9c-db56-42d3-8923-8c5ffdfa637b/homo-sapiens-sapiens

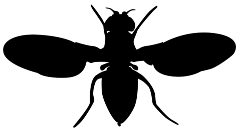

License: CC0 1.0 Universal Public Domain Dedication
Uploaded: 2023 Mar 6 by Maarten van der Sande
Taxon: Drosophila virilis
Link: https://www.phylopic.org/images/07a2b18c-c1f8-4a37-9322-5f901eb567b5/drosophila-virilis

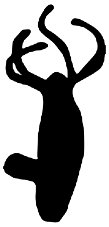

License: CC0 1.0 Universal Public Domain Dedication
Uploaded: 2024 Mar 19 by Tetsuo Kon, MD, PhD
Taxon: Hydra vulgaris
Link: https://www.phylopic.org/images/daaa3a3a-be0b-407f-a4ed-b6175e123a5d/hydra-vulgaris
